# Identifying a cooperative catalytic network for efficient esterase catalysis

**DOI:** 10.64898/2026.05.29.728779

**Authors:** Sharad Sarak, Hong Yang, Colin T. Pierce, Panhavuth Tan, Allison Cafferty, Anne Dao, Dana Junaidi, Ke Shi, Robert L. Evans, Romas J. Kazlauskas

## Abstract

Efficient catalysis is hard to engineer because sequence comparisons reveal only a handful of conserved catalytic residues, and single substitutions rarely speed up catalysis. We hypothesize that efficient catalysis requires a cooperative catalytic network of residues enabling the physical and chemical steps that together determine *k*_cat_. In ancestral hydroxynitrile lyase HNL1, an α/β-hydrolase with poor esterase activity, we identified such a network. Activity-weighted sequence comparison (SigniSite) across ten homologous HNLs and esterases identified 59 positions disfavoring esterase activity, 38 of which differed from esterase EST1. Replacing these 38 with the corresponding EST1 residues, with experimental refinement, yielded a minimal set of fifteen substitutions (HNL1-15) with 64-fold higher *k*_cat_ and ∼400-fold higher *k*_cat_/*K*_M_ than HNL1. Single-substitution reversion in HNL1-16 lowered *k*_cat_ for 15 of the 16 substitutions; activity dropped to ∼2% for twelve reversions and 9-39% for the three that open a tunnel, so each of the 15 contributes with non-additive effects. The crystal structure of HNL1-15 reveals three coordinated structural changes: reshaping the substrate-binding pocket to favor productive ester binding, restoring access to the oxyanion hole, and opening a tunnel for product egress and water entry. These changes arise through backbone rearrangements, altered flexibility and direct active-site contacts. Solvent viscosity kinetics are consistent with isomerization of the enzyme-substrate complex in both fast and slow variants and a chemical step that limits turnover in a slow HNL1 variant. Because cooperativity masked individual contributions, screening on overall rate would have discarded most of the fifteen. Engineering faster enzymes may therefore require multiple simultaneous substitutions to capture cooperativity, and step-specific assays measuring binding, acylation, or product release.

**For Table of Contents Only:** 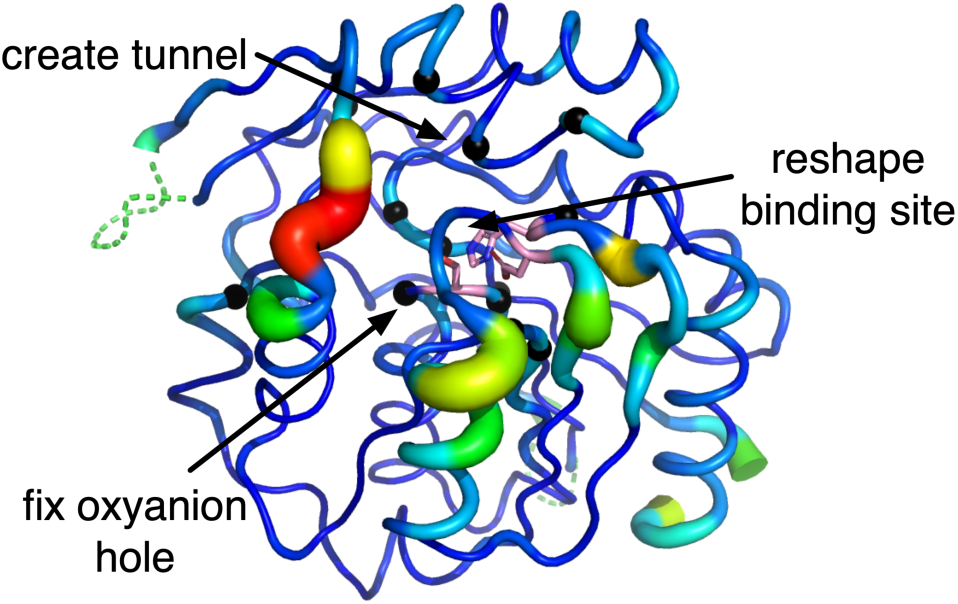

## Introduction

Bioinformatic analysis of enzyme families identifies small sets of residues responsible for catalysis: the catalytic triad of serine hydrolases, the general acid/base in lyases, the metal-coordinating ligands of metalloenzymes^1^. Databases such as M-CSA catalog these chemically indispensable residues across hundreds of enzyme families; substitution of any one abolishes or drastically reduces activity^2,3^.

Directed evolution repeatedly discovers that substitutions far from the active site — in surface loops, at domain interfaces, in second and third coordination shells — substantially enhance or abolish activity^4–8^. Systematic mutagenesis of conserved non-catalytic residues in a class A β-lactamase revealed that second-shell mutations can yield proteins that are stably folded yet catalytically inactive^9^. In *de novo* Kemp eliminases, active-site mutations establish a preorganized center, but distal mutations are additionally required to facilitate substrate binding and product release through conformational dynamics^10^. A residue 12 Å from the catalytic copper in nitrite reductase lowered efficiency ten-fold^11^; one 15 Å from the active site of dihydrofolate reductase lowered hydride transfer 200-fold^12^.

This pattern extends beyond single distal residues to mobile regions that gate access to buried active sites. In cytochrome P450cam, substrate recognition is governed by conformational transitions along the entry channel rather than by residues immediately adjacent to the heme.^13^ In Fe(II)/2-oxoglutarate-dependent oxygenases, flexible lid loops away from the metal center control substrate positioning and gate solvent access, and their sequence varies even among family members sharing the same catalytic machinery.^14^ Most directly relevant, Bridwell-Rabb and coworkers showed that site-selectivity in a Rieske oxygenase is jointly controlled by two active-site residues, three residues lining a substrate-entrance tunnel, and one residue in a flexible loop; transplanting only the tunnel or only the loop reproduces selectivity poorly, and only the full combination transfers it completely.^15^ Active-site engineering is therefore already understood to be commonly insufficient on its own, and cooperative regions beyond the active site have been identified in several enzyme families.^50^ In this work, we identify a cooperative network needed not just to change substrate or positional preference, but to change the chemistry of a reaction from hydroxynitrile lyase to esterase. We find that the cooperative catalytic network involves changes in both chemical and physical steps of catalysis.

In the simplest reactions, *k*_cat_ and *K*_M_ map cleanly onto chemistry and binding. Most enzyme-catalyzed reactions involve more steps, however, so *k*_cat_ and *K*_M_ reflect complex mixtures of elementary steps. For example, serine-hydrolase-catalyzed hydrolysis of *p*-nitrophenyl acetate (*p*N-PAc) involves five steps: substrate binding, formation of an acetyl-enzyme intermediate, release of *p*-nitrophenol and binding of water, hydrolysis of the acetyl-enzyme intermediate, and release of acetic acid, Figure 1. Figure S1 shows the mechanistic steps of this reaction. In an efficient hydrolase like chymotrypsin, physical steps are fast and the chemical step limits the rate. In an inefficient enzyme, the highest barrier may occur at any step, including physical ones such as non-productive binding^16^. *k*_cat_ reflects a purely chemical step only in the simplest cases.

**Figure 1:**
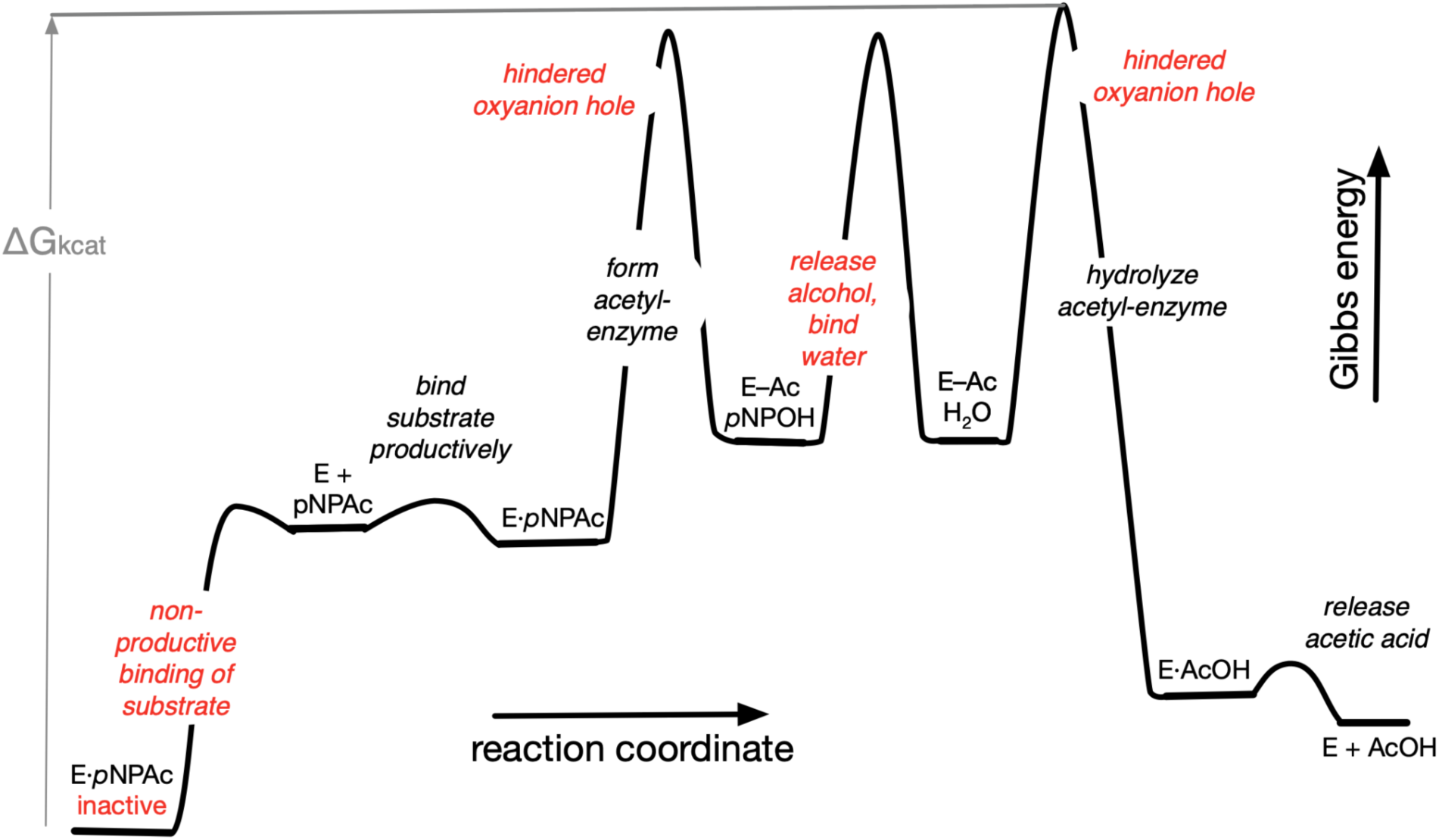
A hypothetical reaction coordinate for *p*-nitrophenyl acetate (*p*-NPAc) hydrolysis catalyzed by an inefficient enzyme. The steps that slow *k*_cat_ (labelled in red) include both physical and chemical steps. Non-productive binding (a physical step) slows *k*_cat_ by stabilizing an inactive form of the E·S complex. A hindered oxyanion hole slows both chemical steps (formation and hydrolysis of the acetyl-enzyme intermediate). Release of *p*-nitrophenol, *p*NPOH, (another physical step) and binding of water slows the sequence between the two chemical steps. The Gibbs energy change that corresponds to *k*_cat_ (ΔG_kcat_) is the distance between the most stable E·S complex (a non-productive form in this example) and the highest transition state along the catalytic path (hydrolysis of the acetyl-enzyme intermediate in this example). Speeding up catalysis, *k*_cat_, requires lowering whichever barrier(s) limit the reaction rate.

Efficient catalysis requires a cooperative catalytic network of two functionally distinct step classes^17^. Chemical steps involve direct covalent bond changes: proton transfer, transition-state stabilization, and positioning of reactive groups. Physical steps create the conditions for chemistry: active-site burial, preorganization of catalytic geometry, and control of substrate access and product release. Neither class is sufficient alone: chemical steps require the physical environment that enables them, and physical steps have no catalytic consequence without the chemical machinery. This obligate interdependence means that neither class of step can be eliminated without abolishing catalysis..

Classifying the steps this way is straightforward; classifying the residues is not. A given residue can contribute to multiple steps. The catalytic histidine is unambiguously chemical, acting as a general acid/base, but its precise positioning depends on a surrounding hydrogen-bond network that is primarily physical. Conversely, a residue that shapes the substrate-binding tunnel may also influence transition-state geometry. We use “chemically essential” and “physically essential” as shorthand for a residue’s dominant contribution, recognizing that most residues participate in more than one way.

This blurring has a practical consequence: bioinformatic methods capture primarily residues with dominant chemical contributions, because chemical essentiality demands exactness - a nucleophile must be a nucleophile - producing the strong conservation that sequence analysis detects. Physically dominant residues are under looser constraint; what matters is shape and flexibility, not the specific residues, so they fall below typical conservation thresholds or are conserved only within narrow subfamilies. Even conserved physical residues are indistinguishable from fold-stabilizing ones by standard sequence alignment^9^. This invisibility explains why the physical side of the catalytic network remains poorly characterized.

The α/β-hydrolase fold superfamily offers an ideal system in which to dissect these contributions. A single Ser-His-Asp triad drives at least 17 distinct reaction types — ester hydrolysis, C–C bond cleavage, epoxide ring opening, dehalogenation, amide cleavage, and others^18,19^ — so reaction type cannot be determined by the triad alone. Differences between a hydroxynitrile lyase and an esterase must arise largely from the physical environment surrounding the triad: the substrate-binding pocket geometry, tunnels that control substrate and product flux, and the degree of active-site burial. Substitutions required to interconvert two α/β-hydrolases sharing the same triad point to dominant physical contributions such as substrate binding, conformational dynamics, and product release. Some of these substitutions may also subtly influence chemical steps, a reminder that the physical/chemical distinction applies most cleanly to steps, not residues.

In previous work, we showed that remodeling the active-site region alone was insufficient to convert *Hb*HNL, hydroxynitrile lyase from rubber tree, into an efficient esterase^17^. Matching *Hb*HNL active-site residues to SABP2 (45% identity) required 14 substitutions plus two stabilizing substitutions. The resulting variant, identical to SABP2 within 6.5 Å of the substrate and encompassing all residues that could directly contact it, was not an efficient esterase. The chemical machinery was in place; physically essential residues beyond the immediate active site were missing.

Second, we demonstrated that adding more distant substitutions created efficient esterases. Extending identity to 10 Å (HNL40, 40 substitutions) and 14 Å (HNL71, 71 substitutions) yielded efficient esterases; HNL71 was approximately twice as active as SABP2 itself^20^. Structural analysis identified three corrective changes: restoring oxyanion hole access, modifying the alcohol-binding region, and creating solvent tunnels. However, the large number of substitutions obscured which substitutions were most important and why.

In this paper, we report a parallel conversion of an ancestral hydroxynitrile lyase, HNL1^21^, to an efficient esterase. Bioinformatic analysis identified 38 residues associated with poor esterase activity. Individual reversion of every candidate established which of these were required, narrowing the set to 15 substitutions (HNL1-15) and confirming that all 15 are individually necessary. This exhaustive test, rather than the identification of one or a few influential distal residues, is what distinguishes this network from prior examples of cooperativity beyond the active site. Structural analysis revealed possible roles for most of the substitutions: one subset reconfigured the substrate-binding pocket to accommodate an ester substrate, while others restored access to the oxyanion hole, created a new tunnel to the active site, and altered the flexibility of the protein. The results demonstrate that cooperative catalytic networks, not the catalytic triad alone, are the functional units of enzyme catalysis, and that engineering new activity requires redesigning the entire cooperative network.

## Results

### Matching residues within the active site does not create an efficient esterase

Esterase activity toward *p*-nitrophenyl acetate (*p*NPAc) was measured spectrophotometrically by following the release of *p*-nitrophenol*/p*-nitrophenolate at 404 nm, and steady-state kinetic constants (*k_c_*_at_, *K*_M_) were obtained by fitting the Michaelis–Menten equation to the resulting initial rates, as detailed in the Experimental Section. Throughout, we treat *k_c_*_at_ as the primary measure of improvement. Raising *k_c_*_at_ increases the reaction rate at every substrate concentration, whereas lowering *K*_M_ increases the rate only below saturation. ^22^ Preparative biocatalysis normally operates at high substrate concentration, so *k_c_*_at_ is the property that limits reactivity. Further, at high substrate concentration, increased affinity for an intermediate can slow the overall rate.^23^ We also report *k_c_*_at_/*K*_M_, which allows comparison with literature values, but the engineering target in this work is *k_c_*_at_..

The x-ray crystal structure of HNL1^24^ contains a glycerol molecule bound in the active site, which is likely retained from the cryoprotectant used during data collection. Since no substrate-or product-bound structure of HNL1 is available, we use this glycerol as a proxy for the substrate-binding site throughout this study. Twenty-three amino acid residues have at least one non-hydrogen atom within 7 Å of this glycerol. A direct contact between heavy atoms occurs at a distance of ∼3 Å, so this permissive 7 Å cutoff includes residues that would contact a differently-shaped substrate or contact a substrate during protein motion. Of these twenty-three residues, eight are identical in HNL1, SABP2 (a modern esterase), and EST1 (an ancestral esterase). We constructed HNL1_all_close_15, which contains fifteen substitutions (T11G, I12A, E79H, C81F, N104T, L106F, F121Y, M122N, L148F, L152F, N156K, F178L, I209A, F210I, K236M) that make all twenty-three active-site residues identical to those in EST1. These substitutions did not improve the *k*_cat_ for *p*-nitrophenyl acetate (*p*NPAc) hydrolysis, Figure 2: 1.6±0.1 min^-1^ for HN-L1_all_close_15 versus 1.7±0.1 min^-1^ for HNL1. The *K*_M_ improved five-fold from 9±1 mM in HNL1 to 1.7±0.3 mM in HNL1_all_close_15. The lack of improvement in *k*_cat_ shows that matching the active-site residues alone does not create an efficient esterase; additional substitutions are also required.

**Figure 2:**
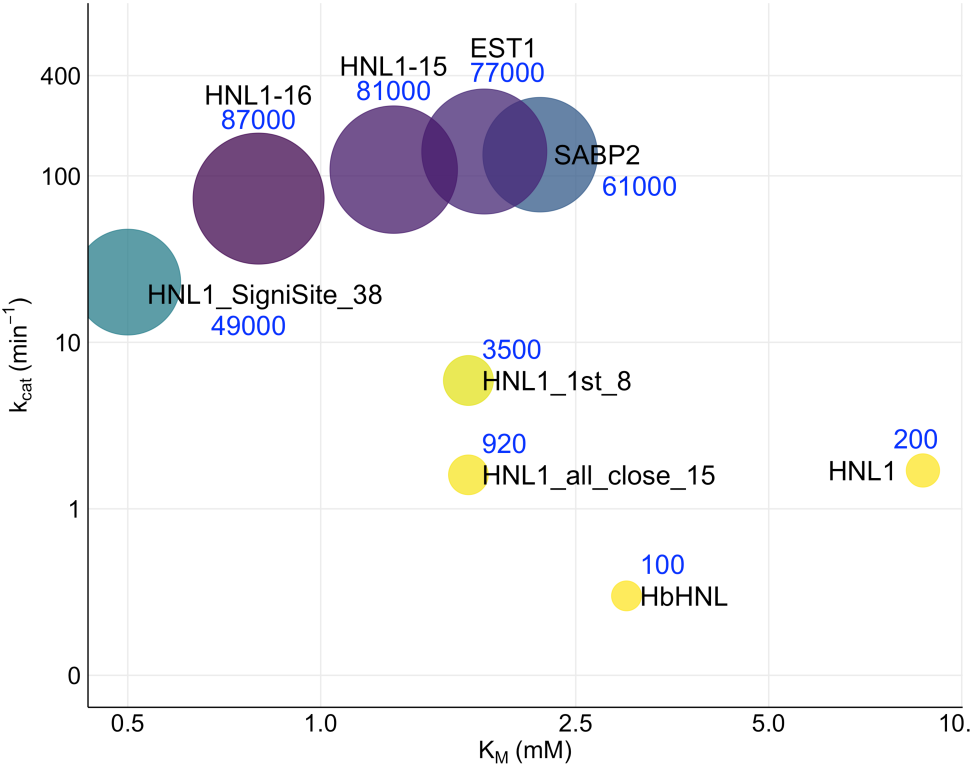
Steady-state kinetic constants for hydrolysis of *p*-nitrophenyl acetate catalyzed by HNL1 variants and selected other enzymes. Catalytic turnover, *k*_cat_, is on the y-axis (logarithmic scale), the Michaelis constant, *K*_M_, is on the x-axis (linear scale). The ball size and color (yellow to blue to purple) indicate the catalytic efficiency, *k*_cat_/*K*_M_ in M^-1^·min^-1^ and the values are shown near the ball in blue. (Table S1 lists the individual values of the kinetic constants.)

### Identifying a set of substitutions that imparts efficient esterase activity in a hydroxynitrile lyase

We used a three-step procedure to identify residues that contribute to esterase activity. First, we selected a small set of homologous HNLs and esterases. Second, we aligned the sequences and ranked residues at each position by their contribution to measured esterase activity, identifying those that disfavor esterase activity. Third, we experimentally tested subsets of substitutions in HNL1 to find a minimum set that imparts efficient esterase activity.

First, we chose ten homologous HNLs and esterases for sequence comparison (Table S2). These included five modern enzymes (two HNLs and three esterases) and five ancestral enzymes (three HNLs and two esterases) reconstructed previously.^21^ The pairwise sequence identity was 57% ±21% between esterases, 78%±9% between hydroxynitrile lyases, and 47%±10% between the two groups. Engineering focused on adding esterase activity to ancestral enzyme HNL1 because it showed high sequence identity (60%) with ancestral esterase EST1. The engineering approach was to replace selected residues in HNL1 with the corresponding residue from EST1. We hypothesized that within this set of closely related enzymes, residues contributing to the physical steps of esterase catalysis would be conserved or mutually compatible.

Second, we identified 38 residues in HNL1 where replacement would increase esterase activity. Using a multiple sequence alignment of the ten enzymes, weighted by measured esterase activity toward methyl pentanoate^21^, we identified residues associated with high esterase activity and low esterase activity at each position in the amino acid sequence, Figure 3. Three esterases (EST1, EST2, SABP2) showed high esterase activity (140-340 min^-1^), two esterases (*Rc*EST, *Rs*EST) showed moderate activity (2.8-7.1 min^-1^), while all five HNLs showed low esterase activity (0.44-1.0 min^-1^) (Table S2). This activity-weighted comparison means that the amino acids in the esterases with moderate activity were weighted lower than those with high activity. The web tool SigniSite^25^ identified 59 positions associated with reduced esterase activity and 61 positions associated with enhanced esterase activity. Of the 59 positions disfavoring esterase activity, 20 were already identical between HNL1 and EST1; ignoring the initial methionine left 38 residues for substitution. These 38 residues in HNL1 were replaced with the corresponding residues in EST1.

**Figure 3.**
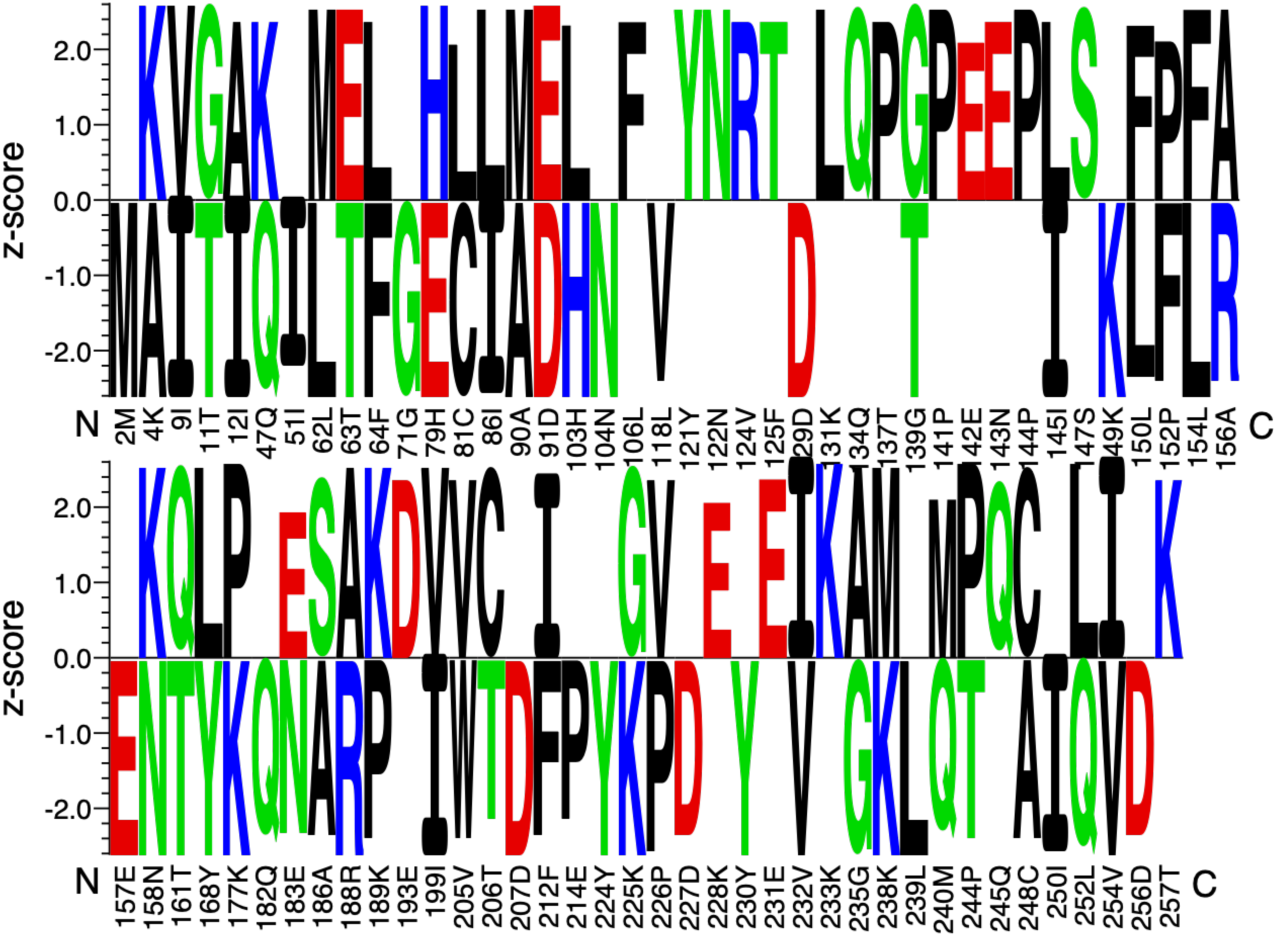
Logo plot of amino acid residues that are predicted to be beneficial (positive scores, 61 residues) or detrimental (negative scores, 59 residues) to esterase activity. The prediction comes from an activity-ranked statistical comparison of ten esterases and hydroxynitrile lyases using SigniSite^20^. To create variant HNL1_signisite_38 we replaced the residues in HNL1 that were predicted to be detrimental to esterase activity with the corresponding residue in esterase EST1. Twenty of the detrimental residues were already absent from HNL1 and the initial methionine was ignored leaving 38 residues for replacement. In most cases, the replacement residue from EST1 matched the beneficial residue in this plot, but if they did not match, the one from EST1 was used. For example, the replacement for C81 was phenylalanine because EST1 contains phenylalanine at this position even though the logo plot predicts leucine as the beneficial residue.

Third, we experimentally tested different subsets of substitutions to find a minimum set that imparts efficient esterase activity. Variant HNL1_SigniSite_38 showed dramatic improvement in esterase activity toward *p*NPAc: a 14-fold improvement in *k*_cat_ from 1.7 to 23 min^-1^, Figure 2 above, Table S1. As expected the hydroxynitrile lyase activity of HNL1_SigniSite_38 toward mandelonitrile dropped significantly. The *k*_cat_ decreased 31-fold from 340±10 min^-1^ for HNL1 to 11±2 min^-1^ for HNL1_SigniSite_38. As described below, further experimentation reduced the number of substitutions to fifteen in variant HNL1-15, which showed even higher esterase activity than HNL1_SigniSite_38: *k*_cat_ = 109 min^-1^, which is comparable to that for SABP2 (134 min^-1^) and for EST1 from which the replacement amino acids were chosen (140 min^-1^).

To reduce the number of substitutions in HNL1_SigniSite_38, we divided them into three groups according to their distance from the bound solvent, glycerol, in the structure of HNL1. The eight 1^st^-shell residues had at least one atom within 7 Å of the glycerol, the eight 2^nd^-shell residues had their closest atom within 7 to 10.5 Å of the glycerol and the remaining twenty-two 3^rd^-shell residues had their closest atom more than 10.5 Å from the bound glycerol (Table S3, Figure 4). The 7 Å cutoff matches the permissive active-site definition used above; the 10.5 Å cutoff separates the next eight nearest residues from the 22 more distant residues. Removing the most distant group (twenty-two 3^rd^ shell substitutions) gave the variant HNL1-16, which showed higher *p*NPAc activity than HNL1_SigniSite_38. The *k*_cat_ increased three-fold to 73 min^-1^ and *k*_cat_/*K*_M_ increased 1.8-fold to 87,000 M^-1^·min^-1^, Figure 2, Table S1. Because removing these 22 positions increased rather than decreased activity, they are not required for the fifteen-substitution network characterized here. We cannot rule out that some of these 22 positions could support a different cooperative network not explored in this study; our reversion analysis establishes that the fifteen retained substitutions are necessary and sufficient, not that no alternative combination could also succeed. Figure S2 shows the sequence alignment of selected enzymes.

**Figure 4.**
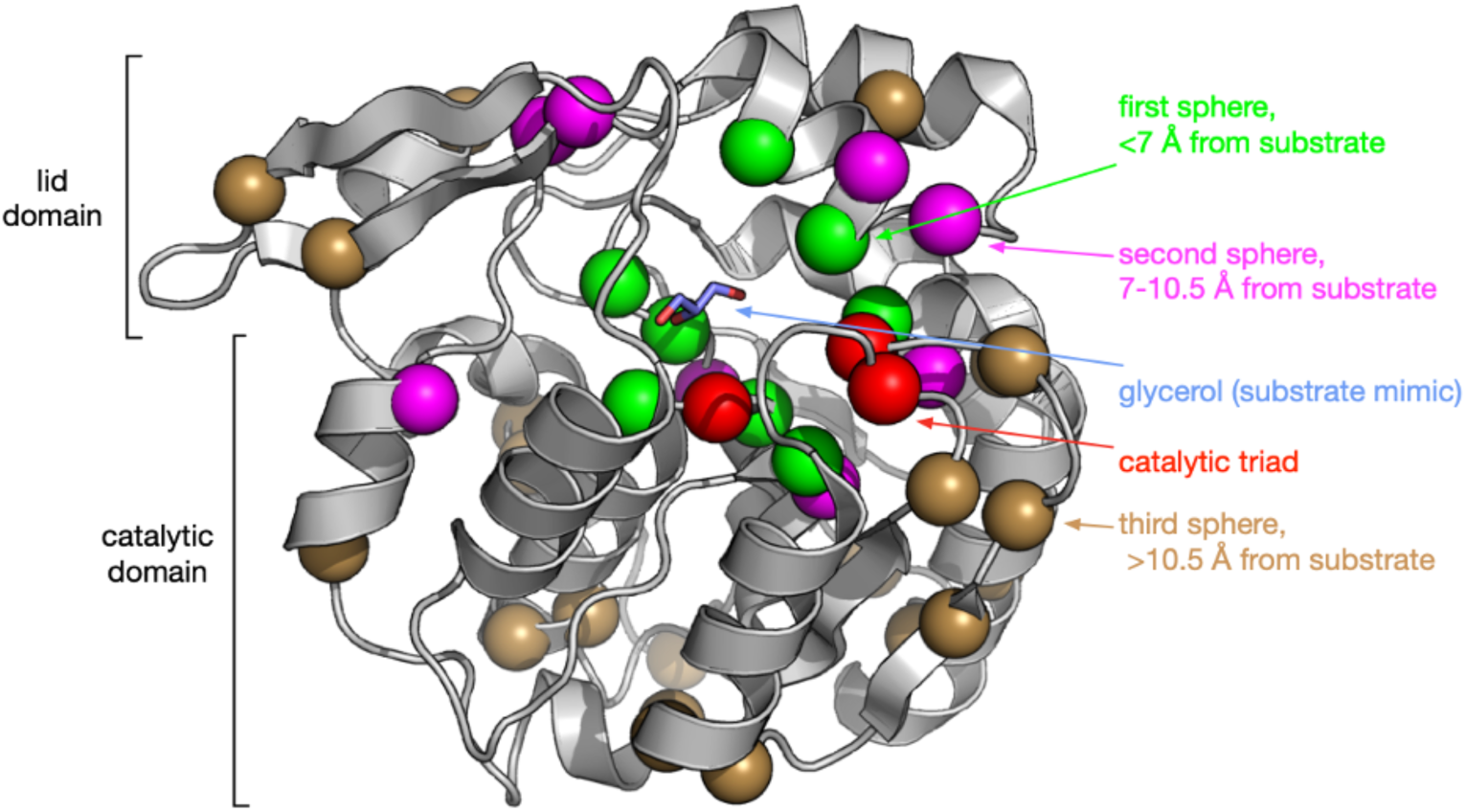
Locations of the thirty-eight substitutions in HNL1_SigniSite_38 shown as spheres at the ɑ-carbon on the x-ray structure of HNL1 (PDB ID 5TDX). The eight first shell residues (green spheres, T11G, I12A, E79H, C81F, N104T, L152F, N156K, K236M) are closest to the active site and have at least one atom <7 Å from the bound glycerol molecule (sticks with pale blue carbons). The eight second shell residues (magenta spheres, I9V, H103L, K147L, E155N, T159Q, K175P, Q180I, L237P) are further away and have at least one atom between 7-10.5 Å from the bound glycerol. The twenty-two third shell residues (brown spheres) are more than 10.5 Å from the bound glycerol and do not significantly contribute to esterase activity.

The 1^st^ and 2^nd^ shell residues were not effective separately; only when combined in HNL1-16 did esterase activity dramatically increase. Variant HNL1_1st_8 (the eight 1st-shell residues defined above) showed improved *p*NPAc activity relative to HNL1, but remained an inefficient esterase. All of these closest eight substitutions, except for L152F and N156K, are in the catalytic domain. The *k*_cat_ of HNL1_1st_8 increased 3.5-fold to *k*_cat_ = 5.9 min^-1^ over HNL1 (*k*_cat_ = 1.7 min^-1^) and the catalytic efficiency increased 18-fold (k_cat_/K_M_ = 3500 M^-1^·min^-1^) over HNL1 (k_cat_/K_M_ = 200 M^-1^·min^-1^). This moderate increase confirms that substitutions within the active site are not enough to create an efficient esterase. Variant HNL1_2nd_8 showed no improvement in *p*NPAc activity relative to HNL1; it contained five substitutions in the lid domain (K147L, E155N, T159Q, K175P, Q180I) and three in the catalytic domain: I9V, H103L, and L237P. The *k*_cat_/*K*_M_ (catalytic efficiency) decreased from 200 in HNL1 to 97 M^-1^·min^-1^ in HNL1_2nd_8. This low value of *k*_cat_/*K*_M_ for HNL1_2nd_8 as compared to SABP2 (61,000 M^-1^·min^-1^) indicates that HN-L1_2nd_8 is a poor esterase. The individual values of *k*_cat_ and *K*_M_ could not be determined because HNL1_2nd_8 showed no saturation with substrate at the highest *p*NPAc concentrations that could be tested. Unlike the 1^st^-shell substitutions, which partially reconfigure the active-site geometry, the 2^nd^-shell substitutions appear to alter the physical environment in ways that are only productive in the context of the reshaped active site. These results establish that the 1^st^-and 2^nd^-shell substitutions form a cooperative unit. The HNL activity of HNL1 toward mandelonitrile (*k*_cat_ = 340 min^-1^) dropped 9-fold in HNL1_2nd_8 (*k*_cat_ of 37 min^-1^), but more sharply, 52-fold, in HNL1_1st_8 (*k*_cat_ of 6.4 min^-1^). These drops show that second sphere residues are also important for hydroxynitrile lyase activity.

The improvement in reactivity (*k*_cat_) of HNL1-16 relative to HNL1 differs for differently shaped alcohols. Esters with large alcohol moieties (*p*-nitrophenol, phenol) improve more (43-and 7.7-fold) than esters with small alcohol moieties (methanol, 2.0-3.2-fold), Table S4. Small alcohols (methanol) may not require reshaping of of the binding site and formation of a tunnel created by the 1^st^/2^nd^ shell substitutions; those structural changes provide less benefit for methyl ester hydrolysis.

### Extensive cooperativity creates efficient catalysis

To assess the contribution of each substitution, we reverted them individually in the HNL1-16 background and measured *p*NPAc hydrolysis (Figure 5, Table S5). Reversion lowered *k*_cat_ for fifteen of the sixteen substitutions. Twelve of these reduced *k*_cat_ to an average of 2% of HNL1-16. The remaining three — L152F, N156K and T159Q — left larger residual activities of 39%, 9% and 17%; these three act together to open the product-release tunnel described below, so each supplies part of a single structural change and reverting one narrows the tunnel rather than closing it. Averaged over all fifteen, reversion dropped the activity to 6% of HNL1-16 activity. The least active variants nevertheless retain substrate binding: *K*_M_ ranges from 0.3 to 2.8 mM, within three-fold of the 0.8 mM measured for HNL1-16, while *k*_cat_ falls as much as 7000-fold (Table S5). The loss of activity therefore reflects a catalytic defect rather than loss of folded protein.

**Figure 5.**
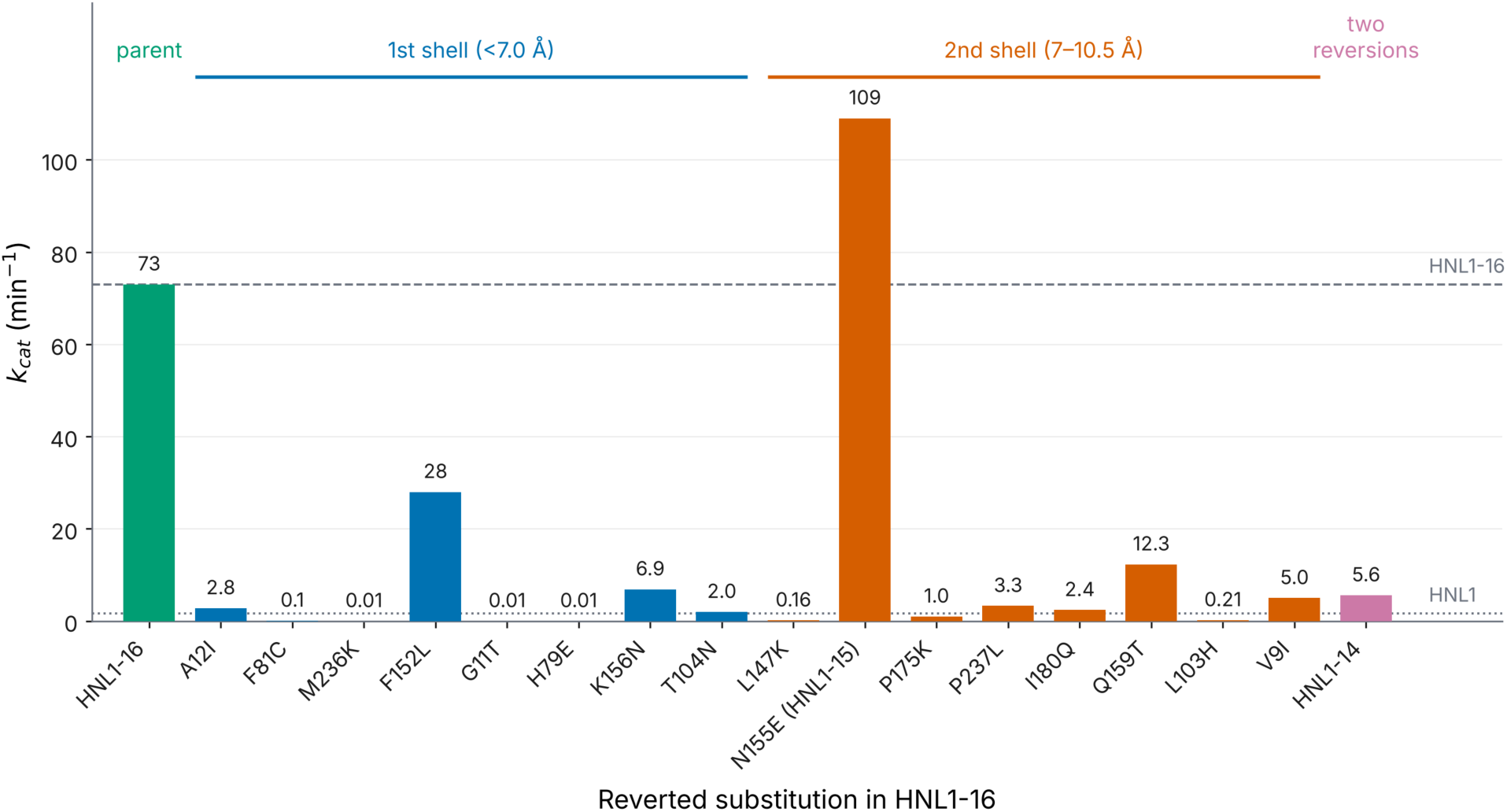
Catalytic activity of HNL1-16 variants in which a single substitution is reverted to the residue found in HNL1. The *k*_cat_ values are given above each bar. Variants are grouped by the distance of the substituted residue from the active site. N155E is the variant designated HNL1-15; HNL1-14 carries two reversions (L103H and T104N). The dashed and dotted lines mark *k*_cat_ for HNL1-16 and for the parent HNL1. Values of *k*_cat_ and *K*_M_ for all variants, with standard errors, are in Table S5.

The sixteenth reversion substitution, E155N, increased esterase *k*_cat_ 1.5-fold to 109 min⁻¹. We designated this variant, which retains the original N155 of HNL1, as HNL1-15, and used it for all subsequent work. HNL1-15 shows no detectable hydroxynitrile lyase activity (<0.1 min^-1^) with mandelonitrile as substrate.

These losses of activity upon reversion are far larger than expected if they acted independently. If the sixteen substitutions in HNL1-16 contributed equally and independently, then each would account for a 1.3-fold increase over the 43-fold total (73/1.7 min⁻¹), and reverting any one would leave about 80% of *k*_cat_. The observed average of 6% is thirteen-fold below that expectation. Even the three tunnel substitutions, whose individual effects are the mildest, fall well short of it. Each substitution therefore contributes to esterase activity, and the contributions are strongly non-additive.

Further evidence for cooperativity comes from testing the first-and second-shell substitutions separately: HNL1_1st_8 and HNL1_2nd_8, containing only the eight first-shell or the eight second-shell substitutions, each gave low activity, demonstrating that neither set alone is sufficient and that substitutions across both shells must act together to produce the full catalytic benefit. Individual pairs also interact non-additively. Reverting L103H alone lowered *k*_cat_ to 0.3% of HNL1-16 (0.21 min⁻¹) and reverting T104N alone to 3% (2.0 min⁻¹). If the two positions contributed independently, reverting both would leave about 0.01% of HNL1-16 activity; instead the double-reversion variant HNL1-14 retained 8% (5.6 min⁻¹), indicating a compensatory interaction between these two positions. Together — the contribution of every substitution within the full set, the failure of either shell subset to recapitulate high activity, and the departure from additivity both across the set and between individual pairs — these results are consistent with the fifteen substitutions forming a cooperative catalytic network. Such cooperativity makes these networks difficult to discover, since the full benefit requires the simultaneous presence of many substitutions.

### Structural basis for improved esterase activity

To identify structural changes caused by the substitutions that make up the cooperative catalytic network, we solved the x-ray crystal structure of HNL1-15 (PDB ID 9DK4; Tables S6-S9) at 1.88 Å resolution and compared it to the structure of HNL1 (PDB ID 5TDX). HNL1-15 shows the same three types of structural changes — reshaping of the active site, tunnel formation, and restoration of access to the oxyanion hole — that accompanied conversion of *Hb*HNL into the efficient esterases HNL40 and HNL71^17^. However, the smaller number of substitutions allows assigning a role to most of them.

#### Binding pNPAc in a catalytically productive orientation

Catalysis begins with a physical step: binding the substrate in a catalytically productive orientation. The substrate-binding sites of SABP2, HNL1 and HNL1-15 differ significantly in shape, Figure 6. SABP2 accommodates both large acid moieties and large alcohol moieties, consistent with its activity toward its natural substrate methyl salicylate (large acid moiety) and toward *p*NPAc (large alcohol moiety), Figure 6A. HNL1, by contrast, accommodates large acid groups, but not large alcohol groups, reflecting the shape of the natural substrate of many hydroxynitrile lyases, mandelonitrile, whose leaving group, cyanide, is small and charged (*k*_cat_ = 340 min^-1^)^18^, Figure 6B. The substitutions in HNL1-15 reshape the binding site to create a hydrophobic space that accommodates the large hydrophobic alcohol moiety of *p*NPAc, Figure 6C.

**Figure 6.**
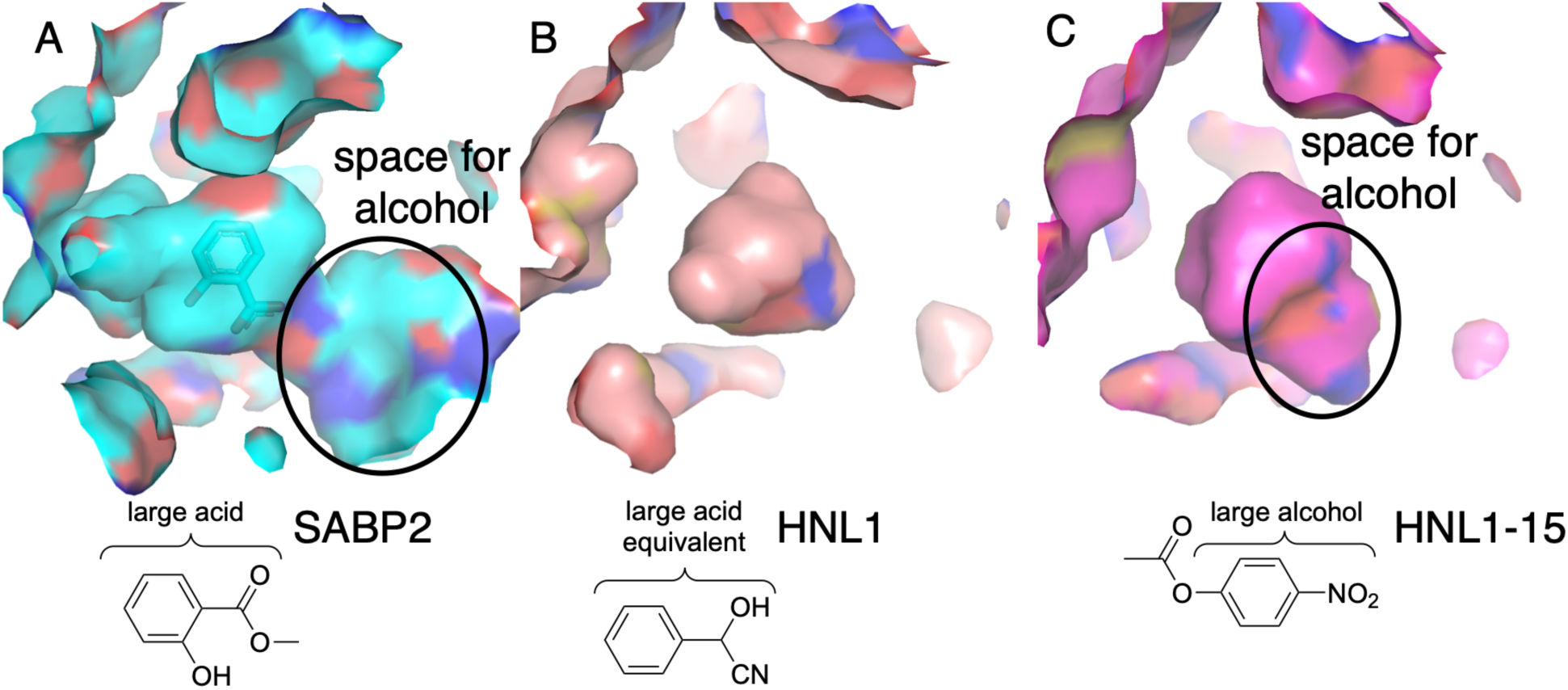
Mandelonitrile cleavage and *p*NPAc hydrolysis require differently-shaped active sites. A) The active site of SABP2 (PDB ID 1Y7I) contains a bound product, salicylic acid (sticks). The active site accommodates both its natural substrate, methyl salicylate, which has a large acid moiety, and *p*NPAc which has a large alcohol moiety. B) The active site of HNL1 does not have space for a large alcohol group because the leaving group on the substrate, mandelonitrile, is small. C) The substitutions in HNL1-15 reshape the active site of HNL1 to create space for a larger alcohol group like that in *p*NPAc.

This reshaping stems from the eight 1^st^-shell substitutions and Ile9Val from the 2^nd^ shell, which also lies nearby. These substitutions collectively remove 144 Å³ (21%) of the original pocket volume while creating 181 Å³ (26%) of new space elsewhere, corresponding to a net 5% (37 Å³) increase in pocket volume: 690 Å³ in HNL1 versus 727 Å³ in HNL1-15, Figure 7A. Beyond volume redistribution, replacing the Glu79–Lys236 salt bridge in HNL1 with His79–Met236 in HNL1-15 makes the substrate pocket more hydrophobic. His79 may additionally form a π–π interaction with the aromatic ring of *p*NPAc, and the Asn104Thr substitution adds a weak hydrogen bond (3.6 Å) to His79 Nε2 that may stabilize its orientation for this π–π interaction.

**Figure 7.**
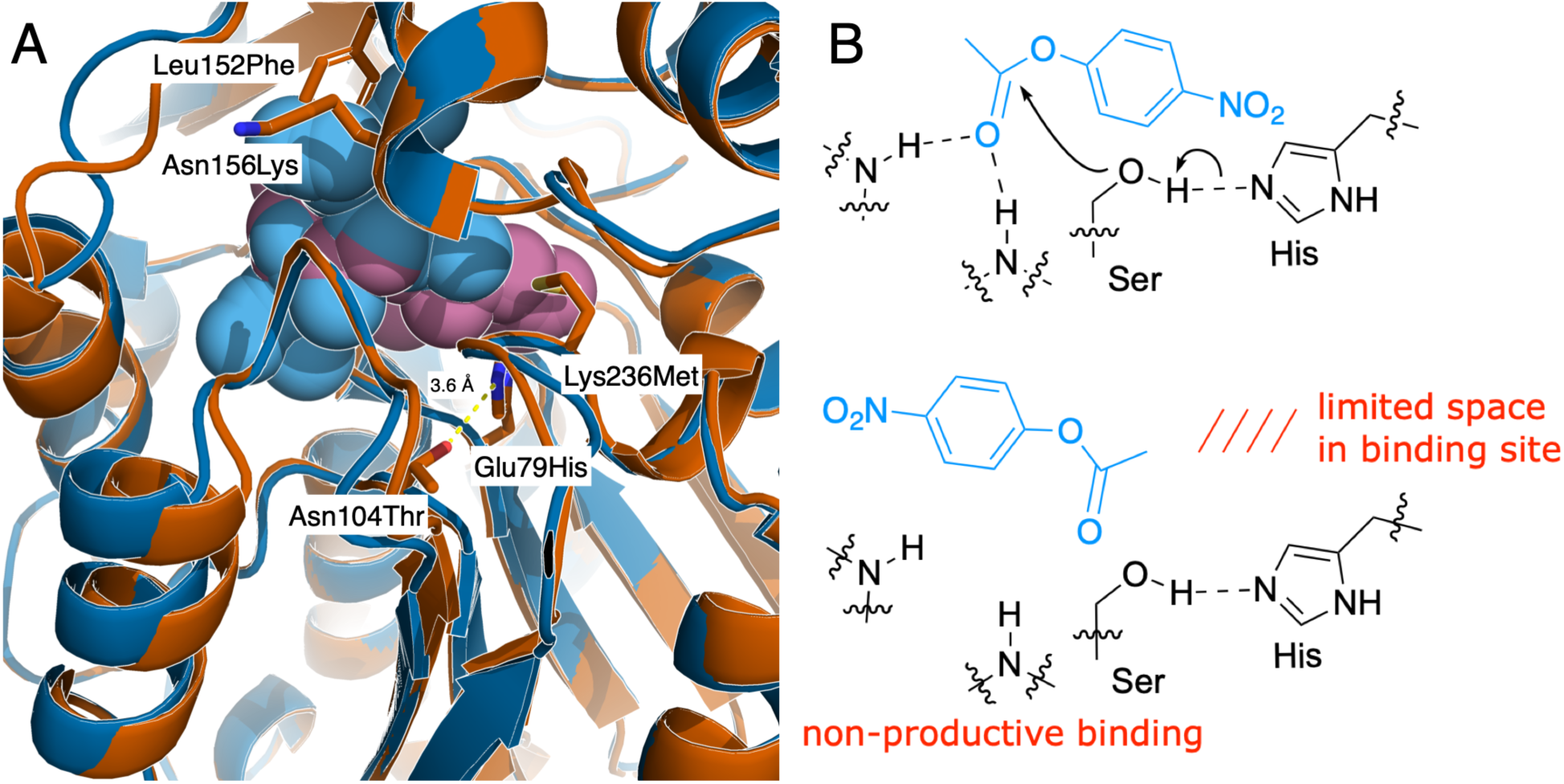
Reshaping the binding site to favor productive binding. A) The eight 1^st^-shell substitutions reshape the active site pocket in HNL1-15 (rose spheres) as compared to HNL1 (blue spheres). Three substitutions are to smaller side chains (Thr11Gly, Ile12Ala, Asn104Thr), one to a similar-sized side chain (Lys236Met), and four to larger side chains (Glu79His, Cys81Phe, Leu152Phe, Asn156Lys). Blue spheres show regions where the pocket of HNL1 is larger, while rose spheres show regions where the pocket of HNL1-15 is larger. The side chains shown as sticks (red-orange carbons) are those in HNL1-15. A dashed line marks a weak hydrogen bond (3.6 Å) between Thr104 Oγ and His79 ε2. Three substitutions (Thr11Gly, Ile12Ala, Cys81Phe) lie on the other side of the pocket and are not visible in this view. B) Productive binding of an ester (top) places the alcohol near the catalytic histidine. If the *p*-nitrophenol moiety does not fit in the binding site, it may bind in a reversed, non-productive orientation (bottom).

Productive binding of *p*NPAc requires positioning the reacting carbonyl carbon near the catalytic serine and the alcohol moiety near the catalytic histidine, which must protonate the leaving group, Figure 7B. Insufficient space for the *p*-nitrophenyl group can force binding in a reversed, nonproductive orientation. Docking calculations (SwissDock^26^) support the hypothesis that reshaping the pocket favors productive binding. Among the top ten docking clusters, SABP2 showed seven productive and three nonproductive orientations; HNL1 showed only three productive, five nonproductive, and two ambiguous orientations. HNL1-15 was intermediate, with five productive, four nonproductive, and one ambiguous orientation. The increase in productive orientations from HNL1 to HNL1-15, though incomplete, is consistent with the reshaping of the binding site partially correcting the steric mismatch that disfavors *p*NPAc binding in HNL1.

#### Formation of the acetyl enzyme intermediate

A chemical step, formation of the acetyl enzyme intermediate, follows the physical step of productive binding of *p*NPAc. A comparison of the catalytic atom positions in SABP2, HNL1 and HNL1-15 reveals one significant change, a change in the side chain rotamer at OX2, Figure 8. In HNL1, the side chain of Cys81 (OX2) points into the oxyanion binding region, which hinders binding of the oxyanion oxygen. In contrast, in SABP2, the side chain of Leu82 (OX2) points away from the oxyanion binding region and does not interfere with the binding of the oxyanion. In HNL1-15, the Cys81Phe substitution at OX2 replaces Cys81 with a different residue than that in SABP2, Phe instead of Leu. Nevertheless, the side chain of Phe81 orients out of the oxyanion binding region, thus enabling the backbone NH of the oxyanion hole to stabilize the transition state. Substitution I12A occupies the oxyanion hole position OX1 and T11G is at the adjacent position. Both of these substitutions also make the oxyanion hole more accessible to the substrate.

**Figure 8.**
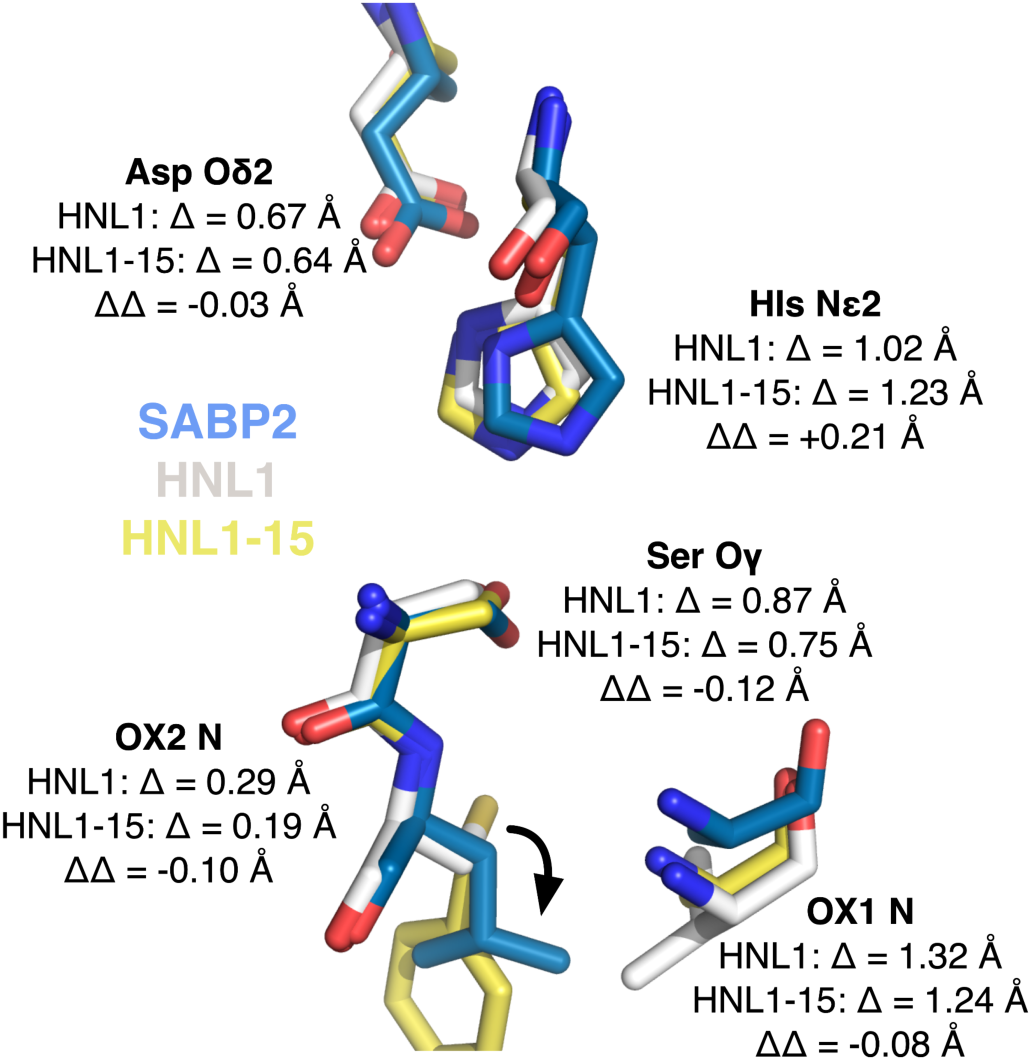
Overlay of the catalytic residues of SABP2 (PDB ID 1Y7H, blue carbons), HNL1 (PDB ID 5TDX, white carbons), and HNL1-15 (PDB ID 9DK4, yellow carbons). The structures of HNL1 and HNL1-15 were overlaid onto the structure of SABP2 using the align function in PyMOL. The HNL1-SABP2 overlay showed an RMSD of 0.60 Å over 207 Cɑ atoms and the HNL1-15-SABP2 overlay showed an RMSD of 0.55 Å over 208 Cɑ atoms. The mean positions of the catalytic atoms (Ser Oγ, His Nε2, Asp Oδ2, OX1 N, OX2 N) in HNL1 differ from those in SABP2 by 0.8±0.4 Å and this distance does not change significantly in HNL1-15 (ΔΔ = 0.02±0.14 Å). The His Nε2 position moves further away, while the other atoms are slightly closer. The largest change was the change in side chain rotamer of OX2 (curved arrow).

Compared to HNL1, the catalytic atoms in HNL1-15 do not move significantly closer to the corresponding positions in SABP2. The mean difference of the catalytic atom positions between SABP2 and HNL1 is 0.83±0.39 Å and between SABP2 and HNL1-15 is 0.81±0.44 Å. This change is not significant. The distance between the His Nε2 increases from 1.02 Å between SABP2 and HNL1 to 1.23 Å between SABP2 and HNL1-15. The other four distances decrease by small amounts.

The p*K*_a_ of the catalytic histidine may increase slightly from 6.5 in HNL1 to 6.9 in HNL1-15 according to PROPKA 3.0^27^ estimates due to the removal of the nearby Glu79-Lys236 salt bridge. Esterase activity was measured at pH 7.2, so this change would be expected to decrease catalytic activity by ∼16%, since more of His235 would be in the protonated, catalytically-inactive, form in HNL1-15. The fact that overall catalytic activity increases suggests that the structural changes dominate the overall effect on catalysis.

#### Release of p-nitrophenol and binding of water

After cleavage of *p*NPAc to the acetyl enzyme intermediate and *p*-nitrophenol, the *p*-nitrophenol must leave and a water molecule must occupy this site. This product release is the second potentially limiting physical step in the catalytic cycle, Figure 9A. The covalent attachment of the acetyl group to the catalytic serine creates a steric hindrance for the product *p*-nitrophenol to leave by the path to the left. An additional path avoiding the acetyl group is needed for the *p*-nitrophenol to leave. The esterase SABP2 contains three large tunnels connecting its buried active site to the solvent, Figure 9B. In contrast, the ancestral hydroxynitrile lyase HNL1 has only one tunnel (center). The 15 substitutions in HNL1-15 create an additional tunnel, making its tunnel architecture resemble that of SABP2. Supporting Table S10 lists calculated tunnel characteristics. The tunnels in both SABP2 and HNL1-15 are too narrow to accommodate entry of the substrate *p*NPAc or exit of the alcohol product *p*-nitrophenol, suggesting that protein motions, such as lid opening, transiently enlarge these pathways.

**Figure 9.**
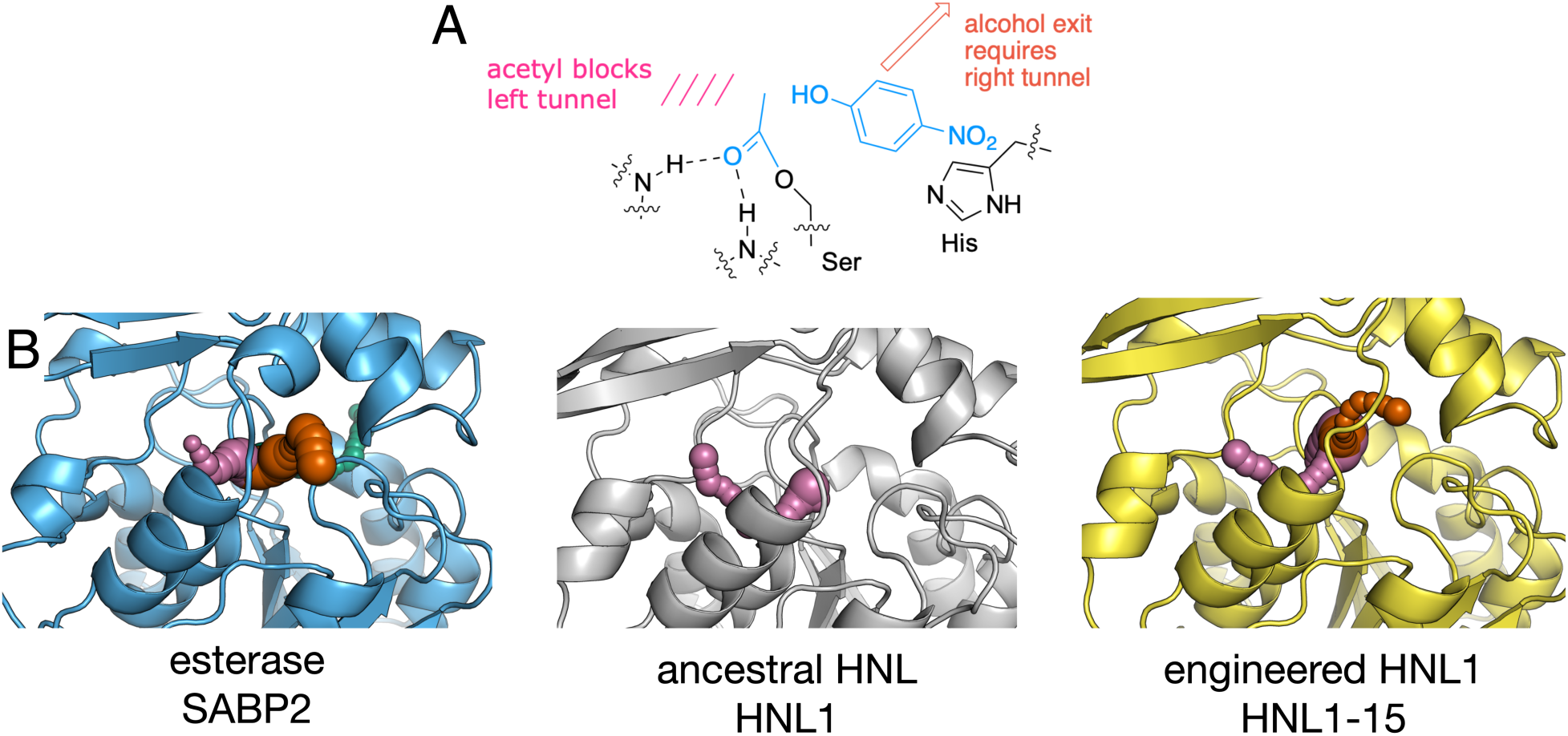
*p*-Nitrophenol release needs a tunnel to the right. A) Formation of the acetyl enzyme intermediate blocks the path to the left; an exit to the right is needed to release the product *p*-nitrophenol and admit water for the reaction to continue. B) Tunnels identified by Caver 3.0^14^. Esterase SABP2 has tunnels to the left (rose) and to the right (vermillion). There is a third tunnel (green) that points to the back. HNL1 has only one tunnel, which points to the left; this lack of a right tunnel likely hinders the exit of *p*-nitrophenol. The substitutions in HNL1-15 create a right tunnel that may allow release of the product *p*-nitrophenol. Tunnel predictions using Caver 3.0 require one protein chain, only natural amino acids, no solvent molecules, no alternative amino acid positions and no missing residues. X-ray structures were modified as needed to fit these requirements as described in the Experimental Section.

The creation of the right tunnel in HNL1-15 involves significant main chain movements. The main chain of HNL1-15 moves in three regions as compared to HNL1, Figure 10A. The largest movement (1.9-3.2 Å) shifts the Cɑ positions along a surface-exposed loop (Phe125-Pro126-As-n127) in the lid domain (residues 108-179) of the protein. The other regions are in the catalytic domain (residues 1–107 and 180–265) and involve smaller shifts, ∼2 Å. One region is in a helix, while the other occurs at residues 205 and 233. Both of these residues are in loops that position the catalytic residues (Asp207, His235). Surprisingly, the shifts in main chain positions do not occur at the substitution sites. The average changes in main chain Cɑ positions in HNL1-15 as compared to the parent HNL1, 0.41±0.40 Å over the 248 residue pairs, is similar to the differences at the 15 substitution sites, 0.45±0.32 Å. The largest Cα shifts occur 3–20 residues from the substitution sites, propagated through secondary structure elements rather than occurring at the substituted positions themselves.

**Figure 10.**
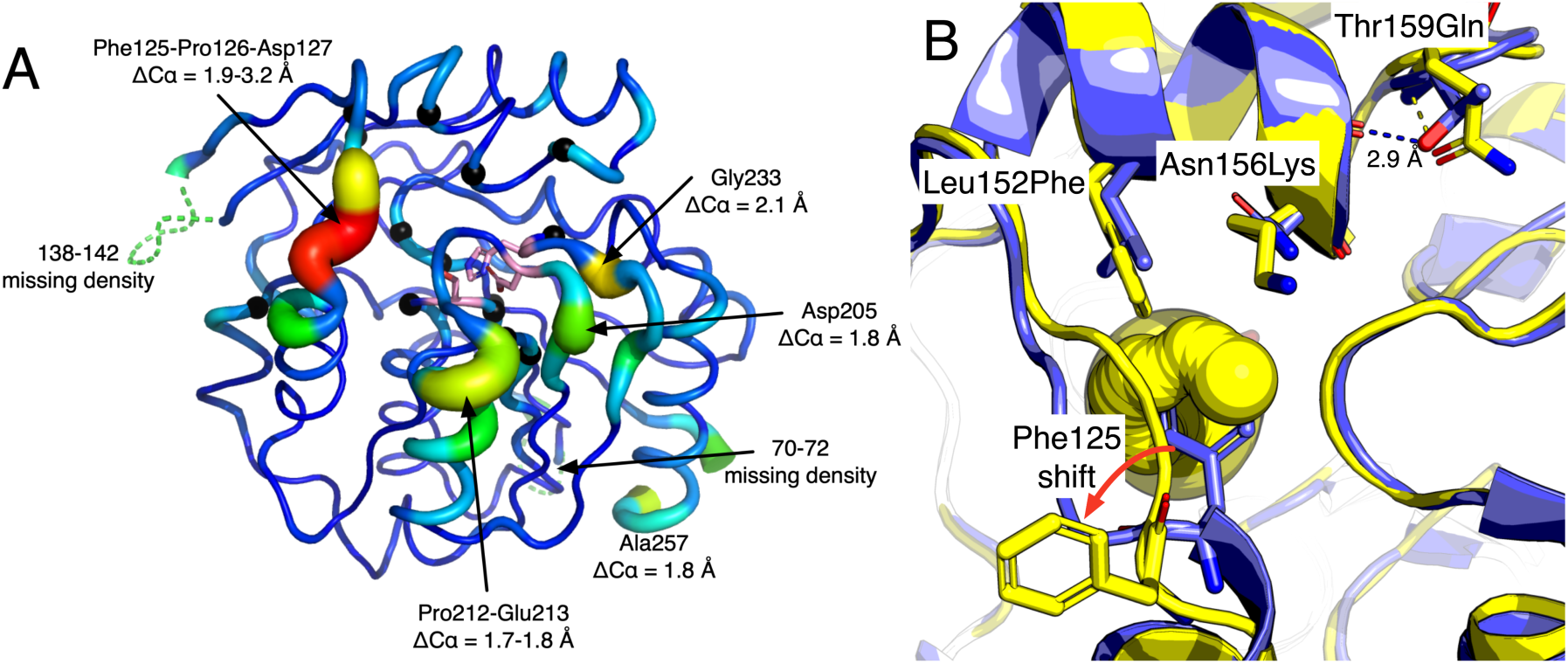
An x-ray structure of HNL1-15 (PDB ID 9DK4) shows shifts in the main chain positions relative to HNL1. A) Differences in the Cɑ positions in HNL1-15 with respect to the parent HNL1. Thicker tubes and warmer colors indicate larger differences in the main chain positions. Dashed lines mark two regions (residues 69-71 and 137-141) where the amino acid positions are not resolved in the structure of HNL1-15. The largest change in Cɑ positions (∼3 Å, red region) occurs at the residues 125-127 and creates a tunnel between the active site and solvent. The catalytic triad is shown as sticks with pink carbon atoms. The changes in Cɑ positions occur not at the 15 substitutions, marked as black spheres at the Cɑ positions, but in nearby regions. B) In HNL1 (blue) Phe125 blocks access to the active site; the view is from the outside looking into the active site. Three substitutions in HNL1-15 (yellow) create a tunnel (yellow) by moving the main chain at Phe125-Pro126-Asn-127. This is the right tunnel (vermillion) in Figure 9. The Leu152Phe and Asn156Lys substitutions on the 149-156 helix increase steric strain causing a shift in the main chain position that pushes Phe125 out of the way. A third substitution nearby, Thr159Gln, breaks a helix-cap hydrogen bond from Thr159 OG to Glu155 O (2.9 Å) that stabilizes this helix.

The substitutions that create this tunnel include both first and second shell substitutions, Figure 10B. Substitutions Leu152Phe and Asn156Lys are in the first shell on a helix, while Thr159Gln in the second shell on a loop following this helix. These substitutions create steric clashes that shift an adjacent loop at positions 125-127 by about 3 Å. This main chain shift creates a new tunnel in HNL1-15. Removing any one of these three substitutions decreases catalytic activity from 100% in HNL1-16 to 39%, 9%, and 17% respectively, Figure 5 above. These decreases (to 22% on average) are less pronounced than at the twelve others in HNL1-15, which show an average decrease to 2%. This less pronounced drop suggests that each residue contributes to forming the tunnel, so that reverting one only narrows the tunnel rather than closing it. Another substitution in this region (Glu155Asn, a second-shell substitution) was initially included in HNL1-16, but was removed to create HNL1-15 because removal increased the catalytic activity by a factor of 1.5.

#### Changes in flexibility

Conversion of HNL1 into the esterase HNL1-15 also altered the main-chain flexibility. The largest changes are two regions with increased backbone flexibility and two with decreased flexibility, Figure S3; none of the fifteen substitutions lie within these regions, but four substitutions lie nearby. Because the two structures were refined independently and differ in resolution and crystal contacts, raw B-factors are not directly comparable. We therefore normalized both structures to a common scale of relative mobility (Z-scores), so that differences reflect whether a residue is more or less mobile than average within its own structure, rather than differing overall B-factor magnitudes.

Both increases in flexibility are associated with substitutions that simultaneously weaken lid–catalytic domain interactions and strengthen lid–lid domain interactions. The increased flexibility of loop-helix-loop 45–52 is most directly explained by K175P and K147L. The K175P substitution breaks two hydrogen bonds from Lys175 Nζ — one to Glu49 Oε2 (a lid–catalytic domain contact) and one to Ser134 Oγ — increasing the conformational freedom of these former partners. The K147L substitution does not break hydrogen bonds but introduces a hydrophobic residue on the surface adjacent to the K175P site, which may contribute to lid–lid packing. The neighboring strand-loop-strand region 134–144 shows propagated flexibility: part of this loop is disordered in the HNL1-15 crystal structure (9DK4) but ordered in HNL1 (5TDX), indicating a substantial increase in conformational freedom in HNL1-15. Although reversion mutagenesis shows that K175P and K147L both contribute to catalysis, Figure 5 above, their precise mechanistic role is not apparent from the structural or flexibility changes alone.

Loop-helix 120–129 lies near the L152F and Q180I substitutions. In HNL1, Leu152 makes van der Waals contact with Trp128 (C–C 3.8 Å); the L152F substitution establishes a stronger aromatic–aromatic interaction with the indole ring of Trp128 (C–C 3.6 Å), strengthening within-lid-domain packing and rigidifying the loop section of 120–129. At the other end of this segment, Q180I removes an H-bond from Gln180 Nε2 to Asp119 Oδ1, breaking a lid–catalytic domain contact. The rigidification of loop-helix 120–129 propagates to the adjacent helix-loop 208–215, which also becomes less flexible in HNL1-15. This loop carries catalytic Asp207, suggesting that L152F and Q180I together reposition and constrain a member of the catalytic triad. Separately, the L237P substitution reduces main-chain flexibility near catalytic His235.

The structural comparison does not suggest a role for substitution H103L despite its being essential for catalysis. Substitution H103L is a stabilizing substitution in *Hb*HNL^31^ and it lies next to N104T, which may orient His79 for interaction with the *p*-nitrophenyl group of the substrate.

The fifteen substitutions are essential for esterase activity in HNL1, but they are not the only combination that can produce an efficient esterase (Table 1, Figure S4). A multiple sequence alignment of 864 SABP2 homologs shows that the amino acids introduced to create HNL1-15 match the family consensus at greater than 90% frequency at only four of the fifteen positions (Val9, Gly11, His79, Met236). The remaining eleven substitutions occur at frequencies ranging from 13% to 78% — no higher than average conservation across all positions — indicating that many alternative amino acid combinations can produce efficient esterases. The oxyanion hole provides a clear example: clearing the oxyanion hole requires a residue bulkier than Cys81 of HNL1, and HNL1-15 accomplishes this with Phe81, but phenylalanine occurs at this position in only 19% of SABP2 homologs. Most (57%) contain leucine; others contain methionine (9%) or tyrosine (9%). At least four different amino acids are compatible with esterase function because the requirement is a bulky side chain, not a specific chemical identity. This functional redundancy contrasts sharply with the catalytic triad, where chemical identity is strictly conserved, and explains why the positions of the cooperative catalytic network do not stand out in sequence comparisons.

**Table 1.**
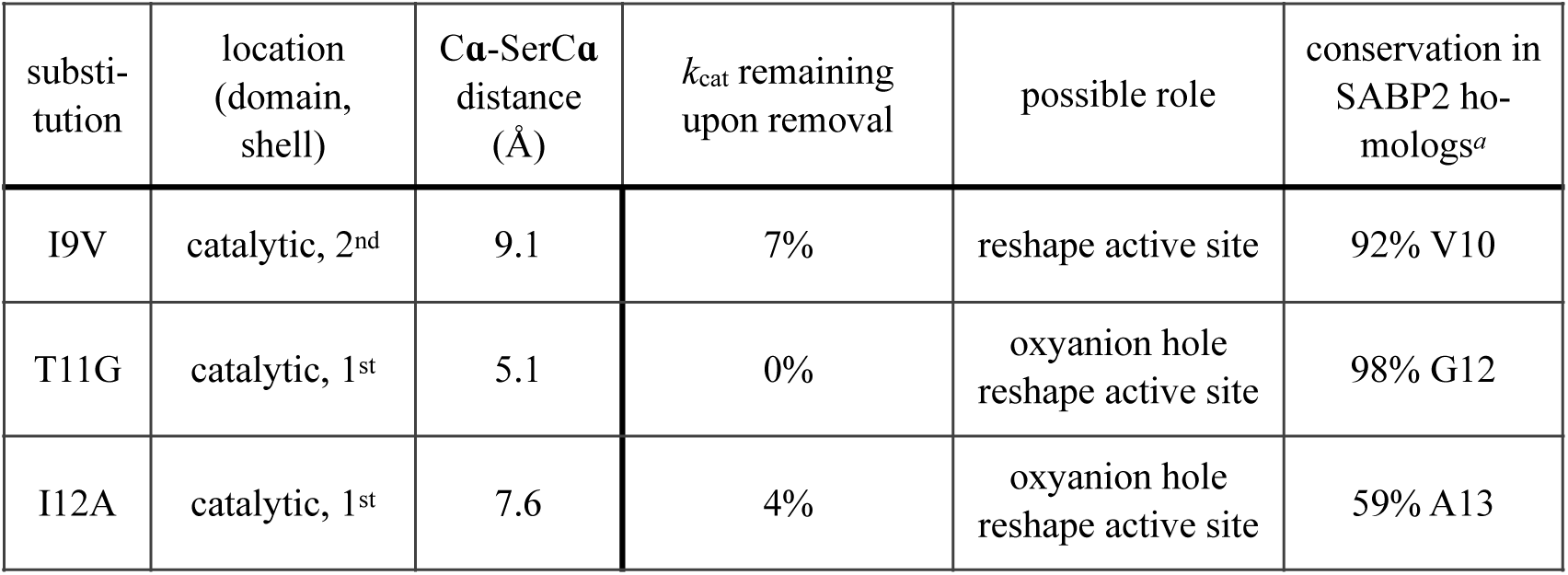
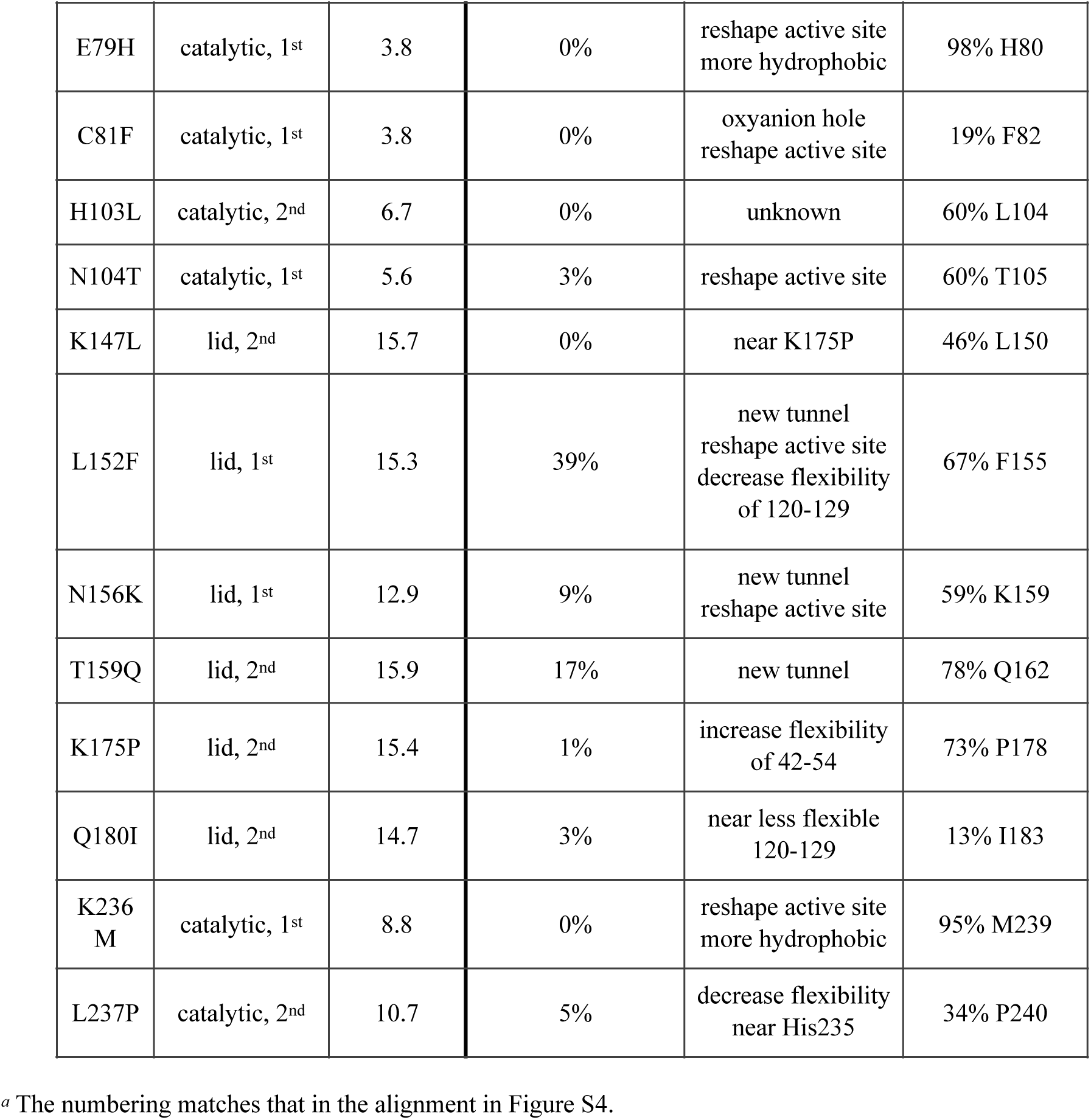
Substitutions in HNL1 to create HNL1-15, their location, the remaining activity upon removing them, their possible role in the catalytic network, and their conservation among homologs of SABP2.

| substi-tution | location (domain, shell) | C**α**-SerC**α** distance (Å) | *k*<sub>cat</sub> remaining upon removal | possible role | conservation in SABP2 ho-mologs<sup>*a*</sup> |
|---|---|---|---|---|---|
| I9V | catalytic, 2<sup>nd</sup> | 9.1 | 7% | reshape active site | 92% V10 |
| T11G | catalytic, 1<sup>st</sup> | 5.1 | 0% | oxyanion hole<br>reshape active site | 98% G12 |
| I12A | catalytic, 1<sup>st</sup> | 7.6 | 4% | oxyanion hole<br>reshape active site | 59% A13 |
| E79H | catalytic, 1<sup>st</sup> | 3.8 | 0% | reshape active site<br>more hydrophobic | 98% H80 |
| C81F | catalytic, 1<sup>st</sup> | 3.8 | 0% | oxyanion hole<br>reshape active site | 19% F82 |
| H103L | catalytic, 2<sup>nd</sup> | 6.7 | 0% | unknown | 60% L104 |
| N104T | catalytic, 1<sup>st</sup> | 5.6 | 3% | reshape active site | 60% T105 |
| K147L | lid, 2<sup>nd</sup> | 15.7 | 0% | near K175P | 46% L150 |
| L152F | lid, 1<sup>st</sup> | 15.3 | 39% | new tunnel<br>reshape active site<br>decrease flexibility of 120-129 | 67% F155 |
| N156K | lid, 1<sup>st</sup> | 12.9 | 9% | new tunnel<br>reshape active site | 59% K159 |
| T159Q | lid, 2<sup>nd</sup> | 15.9 | 17% | new tunnel | 78% Q162 |
| K175P | lid, 2<sup>nd</sup> | 15.4 | 1% | increase flexibility of 42-54 | 73% P178 |
| Q180I | lid, 2<sup>nd</sup> | 14.7 | 3% | near less flexible 120-129 | 13% I183 |
| K236M | catalytic, 1<sup>st</sup> | 8.8 | 0% | reshape active site<br>more hydrophobic | 95% M239 |
| L237P | catalytic, 2<sup>nd</sup> | 10.7 | 5% | decrease flexibility near His235 | 34% P240 |

<sup>*a*</sup> The numbering matches that in the alignment in Figure S4.

To confirm the proposed effects of the observed structural changes in HNL1-15, we measured the kinetic solvent viscosity effects of esterase SABP2 and two HNL1 variants: HNL1_1st_8 (poor esterase activity) and HNL1-15 (good esterase activity). These experiments measured the steady-state kinetics in solutions of differing viscosity, η, due to the addition of sucrose. This analysis detects whether diffusion limits the rate of substrate binding or product release. It also detects solvent-sensitive isomerization of enzyme complexes along the reaction pathway.^28^ The poor esterase HNL1_1st_8 is missing seven catalytically essential residues as compared to HNL1-15. Possible roles of the missing substitutions are changes in flexibility (four substitutions), formation of the alcohol exit tunnel (one substitution), reshaping the active site (one substitution) and unknown (one substitution).

Assuming that acylation of the serine is irreversible in hydrolysis of *p*-nitrophenyl acetate, the variation of *k*_cat_/*K*_M_ with solvent viscosity informs on steps up through formation of the acyl-enzyme (binding, isomerization of enzyme-substrate complex, and acylation). All three enzymes were faster in more viscous solutions, known as an inverse hyperbolic pattern, Figure 11. The ratio (*k*_cat_/*K*_M_)_o_/(*k*_cat_/*K*_M_)_η_ decreased as the viscosity increased. This behavior is consistent with a solvent-sensitive isomerization of the enzyme-substrate complex. The more viscous solutions favor the downstream species in the catalytic pathway, thus increasing *k*_cat_/*K*_M_. One possible mechanism is the formation of a non-productive enzyme-substrate complex, which equilibrates with a catalytically productive enzyme-substrate complex. Higher viscosity favors the productive enzyme-substrate complex, so *k*_cat_/*K*_M_ increases at higher solution viscosity.

**Figure 11.**
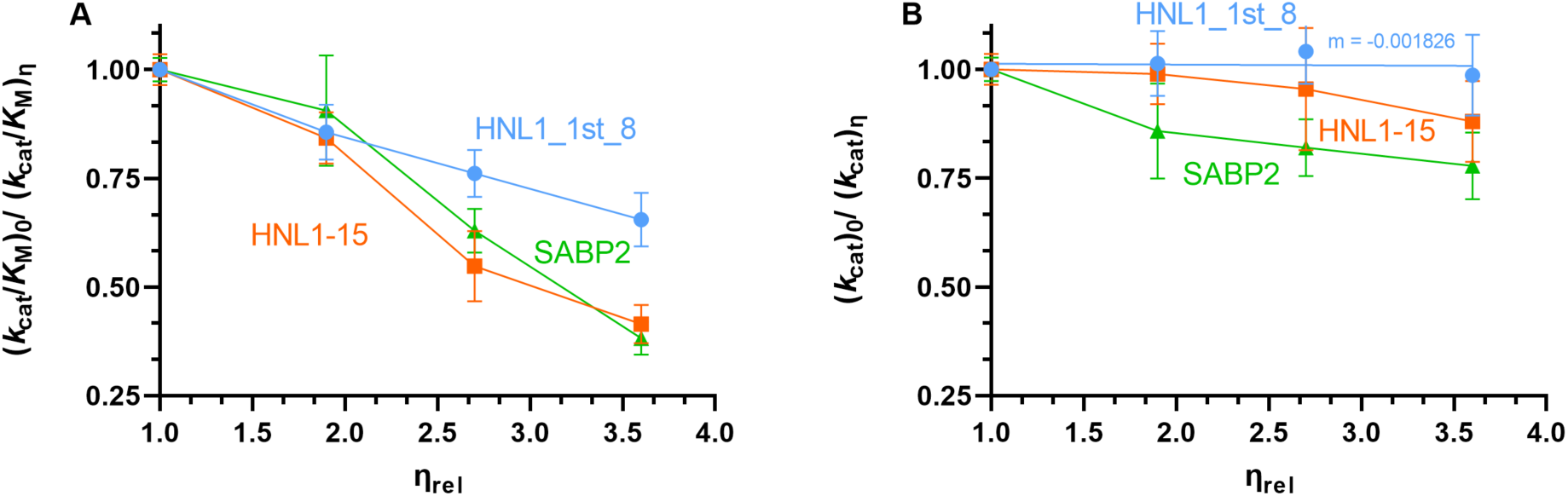
The viscosity dependence of the esterase activity toward *p*-nitrophenyl acetate for esterase SABP2 and two HNL1 variants: HNL1_1st_8 (poor esterase activity) and HNL1-15 (good esterase activity). A) All three enzymes increased *k*_cat_/*K*_M_ in more viscous solutions, which is consistent with a solvent-sensitive isomerization of the enzyme-substrate complex. B) The *k*_cat_ for HNL1_1st_8 does not vary with solvent viscosity, which is consistent with a rate-limiting chemical step. In contrast, SABP2 and HNL1-15 show an increase in *k*_cat_ with increasing solvent viscosity, which is consistent with fast steps downstream of acylation. For this reason the viscosity-sensitive isomerization of the enzyme-substrate complex remains visible in the *k*_cat_ dependence. The line for HNL1_1st_8 in panel B is a best fit line. All other lines connect the data points and serve only to guide the eye.

The variation of *k*_cat_ with solvent viscosity reports on all steps downstream of formation of the acyl enzyme intermediate: dissociation of *p*-nitrophenol from the acyl-enzyme·*p*-nitrophenol complex, the deacylation chemical step, and release of acetate. The solvent viscosity dependence of *k*_cat_ for HNL1_1st_8 shows a slope of zero, that is, no change as solvent viscosity increases, Figure 11B. A dissociation step, such as *p*-nitrophenol release from the acyl-enzyme complex, would be expected to slow in more viscous solutions approaching a slope of 1 as it becomes rate-limiting. The observed zero slope points to deacylation, a viscosity-independent chemical step, as rate-limiting for HNL1_1st_8.

For SABP2 and HNL1-15, the inverse hyperbolic pattern seen for *k*_cat_/*K*_M_ persists for the viscosity dependence of *k*_cat_, Figure 11B. This persistence indicates that the downstream steps (*p*-nitro-phenol dissociation, deacylation, acetate release) are kinetically invisible and the enzyme-substrate-isomerization signature carries through. Thus, the chemical steps and product release are fast in SABP2 and HNL1-15, as suggested by the presence of a tunnel and other structural features essential to catalysis. In contrast, a chemical step, deacylation, limits the rate of HN-L1_1st_8.

This experiment also suggests how to further increase the esterase activity of the good esterases in the future. Catalysis by SABP2 and HNL1-15 is limited by the isomerization of an enzyme-substrate complex. Shifting this equilibrium toward the productive complex by eliminating non-productive binding modes would remove this bottleneck and increase both *k*_cat_/*K*_M_ and *k*_cat_.

## Discussion

The three categories of structural change in HNL1-15 each address a distinct step in the catalytic sequence. Repairing the oxyanion hole (Thr11Gly, Ile12Ala, Cys81Phe) accelerates the chemical steps — acylation and deacylation — by stabilizing the developing negative charge on the carbonyl oxygen in the tetrahedral intermediates. Reshaping the substrate binding site with the eight 1^st^-shell substitutions plus I9V accelerates reaction by correctly orienting the ester for nucleophilic attack by the catalytic serine. Creating a product release tunnel with three substitutions facilitates departure of the *p*-nitrophenol leaving group and entry of the catalytic water that hydrolyzes the acyl-enzyme intermediate. Active site substitutions alone produced only modest gains; when physical steps collectively limit the rate, improving chemistry alone cannot produce a large rate increase. The full rate enhancement emerges only when all categories of barrier are lowered simultaneously, consistent with the strong cooperativity observed among the fifteen substitutions. These fifteen substitutions define one sufficient cooperative network, not necessarily the unique one.

Our procedure for finding residues outside the active site that contribute to catalysis involved several arbitrary choices — the number and selection of sequences used for comparison, the ester substrate chosen to rank catalytic activity, the target enzyme chosen for engineering and the choice of replacement residues. More than half of the 38 predicted substitutions (23 out of 38, 61%) were false positives since removing them to create HNL1-15 maintained and even increased esterase activity. Whether this approach would succeed more broadly — for other enzyme families, other activity transitions, or with a different sequence set — remains to be tested.

Other tools that might identify catalytically important residues include correlated mutation network analysis such as the CorNet tool,^29^ which identified substitutions in an esterase that increased reaction rate and enantioselectivity.^30^

Notably, the fifteen substitutions that compose the functional network all lie within 10.5 Å of the active-site glycerol, which is closer than the 12–20 Å distances reported for individually influential residues in other enzymes. This closeness does not diminish their significance: seven of them lie outside the primary substrate-contact shell (>7 Å), are structurally and functionally distinct from the catalytic triad, and would be missed by an active-site-only engineering strategy. It does indicate that, for this conversion, the physically essential network is compact and contiguous with the active site rather than dispersed across the fold. Whether the spatial extent of such networks scales with the magnitude of the required functional change — a modest activity increase versus the reaction-type change reported here — is a question this result raises but a single example cannot answer.

The specific examples from HNL1-15 illustrate why residue classification is inherently imperfect. Val9 and Gly11 are conserved (>90% in homologs) and reshape the substrate-binding pocket, contributing to binding. Gly11 also restores access to the oxyanion hole, thus contributing to catalysis. Phe81 also restores access to the oxyanion hole but is not conserved, and would be missed by a conservation-based classification scheme. His79 and Met236 (also >90% conserved) eliminate the Glu79-Lys236 salt bridge, increasing active-site hydrophobicity; this change simultaneously promotes binding of the nonpolar substrate *p*NPAc and alters the active-site electric field in ways that could stabilize bond making and breaking. The step-level distinction between chemical and physical catalysis is clean, but individual residues rarely respect it.

The network is better understood as a set of property requirements — shape, electrostatics, hydrophobicity — than as a list of specific amino acids or positions. Sequence alignments of homologs show that 11 of the 15 positions tolerate multiple residues: Phe81, for example, can be replaced by Leu, Met, or Tyr, indicating that the requirement is for a bulky hydrophobic residue at this position, not phenylalanine specifically. This degeneracy at the amino acid level likely extends to the positional level as well: a product release tunnel, for instance, might be created through substitutions at alternative sets of positions. The many natural esterases within the α/β-hydrolase fold — each with a distinct active-site architecture around the same catalytic triad — confirm that multiple sequence solutions satisfy the same functional requirements. Physical-role residues therefore evade conservation analysis: the sequence-level solution space is large even when the property-level requirement is strict.

Physical rather than chemical steps limit reaction rates in several well-studied enzymes, including triosephosphate isomerase, adenylate kinase, and dihydrofolate reductase^31–34^. Tunnels can also limit the overall reaction rates^17,35,36^. Distant mutations that improve catalysis through physical effects appear in several enzymes—directed evolution of a Kemp eliminase reshaped the conformational landscape to increase the population of catalytically competent states^7^; distal mutations in LovD altered active-site dynamics through an allosteric network without changing active-site chemistry^37^; point mutations distant from the active site of chorismate mutase altered electronic structure and dynamics in ways that affected catalytic rates^38^. Lassila and coworkers^39^ demonstrated that many amino acids are tolerated at non-catalytic positions in chorismate mutase.

The structural changes that enable esterase activity disrupt lyase activity. HNL1-15 has no detectable lyase activity, confirming that the esterase-enabling changes are incompatible with the lyase mechanism. In a serine esterase, the oxyanion hole stabilizes the developing negative charge on the carbonyl oxygen during formation of the tetrahedral intermediate^40^, Figure S1. In HNL1, the Thr11 and Cys81 side chains block the oxyanion hole and are essential for lyase activity^41^. This blockade prevents the cyanohydrin hydroxyl from binding in the oxyanion site, which would create a dead-end orientation incompatible with the elimination mechanism.

Reshaping the binding site presents a similar conflict. During ester hydrolysis, the *p*-nitrophenyl leaving group occupies the alcohol-binding region of the active site, whereas during cyanohydrin cleavage the phenyl group of mandelonitrile occupies the acyl-binding region—effectively the opposite orientation within the same pocket^21,42^. The substitutions in HNL1-15 that reconfigure the pocket to position the ester for nucleophilic attack by the catalytic serine simultaneously mis-align mandelonitrile, whose geometry requirements differ substantially from those of *p*NPAc.

Both reactions require active-site desolvation^43^, but achieve it differently. Cyanohydrin cleavage is a concerted chemical step: the substrate enters, the enzyme cleaves the cyanohydrin bond, and the products exit by the same path. Additional tunnels would admit water, weakening the electro-static interactions the lyase depends on. In the esterase, the departing leaving group may transiently occlude the tunnel, potentially limiting bulk solvent access during the chemical steps. In the lyase, the absence of tunnels maintains desolvation.

### Epistasis mediated through catalysis as well as protein structure

Epistasis (cooperativity) can be specific, arising from direct or indirect physical interactions, or nonspecific, arising from nonlinear mapping between a physical property, such as stability, and biological function^44^. In HNL1-15, structural coupling is certainly present: the fifteen substitutions collectively reshape the active site, and a residue that opens a substrate tunnel necessarily alters the environment in which binding-site residues operate. However, structural coupling alone may not fully account for the observed cooperativity. Residues responsible for oxyanion hole stabilization, substrate orientation, and product egress contribute to distinct steps along the catalytic trajectory, and co-operativity among them arises in part because catalysis is a serial process — only when all barriers are lowered simultaneously does the full rate enhancement emerge. Distinguishing these contributions experimentally would require pre-steady-state kinetics to assign each substitution to the step it primarily affects — beyond the scope of this work.

Recent work on other enzymes supports the notion that the catalytic process itself generates non-additivity. Consistent with this, Buda & Tokuriki^45^ recently showed using kinetic modeling that the multi-step nature of catalytic cycles inherently generates non-additivity in *k*_cat_ and *K*_M_. Petrović *et al.*^46^ similarly argued that conformational dynamics — themselves a physical step — are integral to catalysis across enzyme families, making process-level cooperativity a general expectation rather than an exception. In a directed evolution study of β-lactamase OXA-48, Fröhlich et al.^47^ found that four mutations increased antibiotic resistance 40-fold despite individual effects of twofold or less; the synergy arose because successive mutations removed sequential rate-limiting barriers rather than interacting physically. Judge et al.^48^ observed substrate-dependent pairwise cooperativity between active-site residues in CTX-M β-lactamase, consistent with epistasis mediated through the catalytic cycle rather than through fixed structural contacts. The homologous isopropylmalate dehydrogenases from *E. coli* and *Pseudomonas aeruginosa* differ at 168 of 365 positions; exchange of single *E. coli* residues with their *Pseudomonas* counterparts reduced activity for 63 of those positions, which scatter throughout the structure with only one contacting the substrate directly^49^. These cryptically important residues are consistent with process-level epistasis: their cooperativity reflects contributions to distinct catalytic steps — cofactor binding, loop closure, product release — rather than direct structural interaction. This framework predicts that any enzyme with a buried active site and multi-step mechanism — the common case — should show analogous process-level cooperativity.

## Conclusions

Converting ancestral hydroxynitrile lyase HNL1 into an efficient esterase required fifteen substitutions that function as a single cooperative unit: eight within the substrate-contact shell and seven beyond it. Neither group suffices alone. Making all twenty-three active-site residues identical to those of a proficient esterase improved *K*_M_ five-fold but left *k*_cat_ unchanged, and the second-shell substitutions by themselves slightly reduced activity. Together they raise *k*_cat_ sixty-four fold. The first-shell substitutions reshape the binding pocket and clear the oxyanion hole through direct contact with the substrate; the second-shell substitutions act at a distance, shifting the back-bone and altering flexibility to open a product-release tunnel and constrain the loops that position the catalytic triad. Neither change is the work of a single substitution: three substitutions act together to shift Phe125 and open the tunnel, and several reshape the pocket. A cooperative catalytic network is not a set of remote residues acting in place of active-site ones — it spans both shells, and the contributions are interdependent.

Such networks evade conservation-based detection because their requirements are physical rather than chemical. Position 81 needs a bulky side chain, a requirement met by Phe, Leu, Met, or Tyr; eleven of the fifteen positions tolerate multiple residues in homologs. A network specified by properties rather than identities leaves little sequence signal. The fifteen substitutions reported here are one sufficient solution, but not the only one.

These results bear on a widespread assumption: that *k*_cat_ reports the chemistry of catalysis, and that raising it is a matter of better transition-state stabilization. It is not. *k*_cat_ is set by whichever step limits turnover — productive reorientation of the bound substrate, transit of product through a tunnel, entry of the catalytic water, or chemistry itself. Stabilizing the transition state raises *k*_cat_ only when a chemical step is rate-limiting; otherwise it lowers a barrier that was not the tallest and the overall rate does not change. The limitation also moves as engineering proceeds: a chemical step limits turnover in a variant lacking the 2^nd^ shell substitutions, whereas a physical step, productive substrate binding, limits HNL1-15.

Beneficial substitutions are hard to detect because cooperativity operates at two levels, and they compound. Several substitutions may be required to accomplish a single structural change: three act together to open the product-release tunnel, and reverting any one of them in the HNL1-16 background leaves 9–39% of activity — a milder loss than the ∼2% remaining after reverting any of the other twelve. That difference is the signature of a shared job. Each of the three contributes part of the backbone shift that moves Phe125, so losing one narrows the tunnel rather than closing it, whereas the other twelve each carry a change no other substitution supplies. Separately, improving one step yields little while other steps remain limiting. A substitution therefore appears neutral for either of two reasons: its partners in the same structural change are absent, or the step it improves is not the one setting the rate. In both cases it is discarded. Screening on overall rate alone would have eliminated most of the fifteen substitutions reported here.

Step-specific measurements address the second problem: measuring productive binding, acylation, and product release directly shows whether engineering has improved a step, whether or not that step currently limits the rate. They do not address the first. A substitution contributing to a change that requires two others will register as neutral in any assay until its partners are present, so step-specific criteria must be paired with testing substitutions in combination rather than singly. An enzyme is no faster than its slowest step, and that step is often not a chemical one.

## Experimental Section

### Protein production and purification

The genes for HNL1_all_close_15, HNL1-16, HN-L1_1st_8, HNL1_2nd_8, HNL1-15, and all single-reversion variants were codon-optimized, synthesized, and cloned into pET21a(+) with a C-terminal His6-tag and linker by Twist Biosciences and verified by DNA sequencing. All amino acid sequences are in the supporting information and Table S6 includes the DNA sequence for HNL1-15. The proteins were expressed in *Escherichia coli* BL21 as described previously for related proteins^20^. All proteins were purified by nickel affinity chromatography. The HNL1-15 protein for x-ray crystallography was further purified by size exclusion chromatography and yielded a final protein concentration of 5 mg mL^-1^.

### Crystallization of HNL1-15

Initial crystallization screening was performed robotically as described previously^20^ using the FUSION protein crystallization screen^52^ assembled from components of the MORPHEUS screens^53^ from MiTiGen (Ithaca, NY). Sitting drop vapor diffusion trials contained 0.1 µL protein sample (5 mg mL^-1^ protein) and 0.1 µL of reservoir solution. Crystals appeared within 1 day in C8, which contained 5% (w/v) PEG 20,000, 25% (w/v) 1,1,1-tris(hydroxymethyl)propane, 1% (w/v) NDSB 195 (dimethylethylammonium propane sulfonate), 1 mM of each alkali (barium acetate, cesium acetate, rubidium chloride, strontium acetate) and 0.1 M *N,N*-bis(2-hydroxyethyl)-2-aminoethanesulfonic acid/triethylamine pH 7.5 and grew to the full size of 50 x 80 x 30 µm within 3 days, Table S7.

### Structure determination and refinement

Diffraction data were collected at National Synchrotron Light Source II, Brookhaven National Laboratory, AMX beam line, to a resolution of 1.88 Å, Table S8. Initial phases were obtained via molecular replacement with Phaser in the Phenix suite^54^, using the parent HNL1 structure (PDB ID: 5TDX) as the search model. Molecular replacement gave a log-likelihood gain (LLG) of 5148 and a translation-function Z-score of 65.8. The refinement was based on 73,413 reflections of which 4794 reflections (5%) were excluded for calculation of *R*_free_. Initial refinement statistics were improved with PDB-REDO^55^, which reduced the initial *R*_work_ from 0.3516 to 0.2304 and *R*_free_ from 0.3927 to 0.2750. Subsequent manual rebuilding and refinement over 67 rounds in total yielded a final structure with an *R*_work_ of 0.1837 and an *R*_free_ of 0.2283, Table S9. The coordinates and structure factors were deposited in the Protein Data Bank under accession code 9DK4.

The asymmetric unit contains four protein chains. The N-terminal Met1 and C-terminal His tag and associated linker 258-272 are disordered (not modeled) in all four chains. In addition, there are two disordered loops. Loop 70-72 is disordered (not modeled) in chains A, C, and D, but chain B contains a modeled Glu72. Loop 137–142 (TSNNET) varies most between chains. Chain B is the most ordered (only 139–141 missing). Thr137 is disordered in chains C and D, and Thr142 in chains A and C. Unassigned electron density was observed near residue 11 in all four chains and near residue 52 in chains C and D. None of the crystallization components provided a satisfactory fit, so this density was left unmodeled.

### Esterase activity

Esterase activity was measured at room temperature (∼22 °C) by monitoring the hydrolysis of *p*-nitrophenyl acetate (*p*NPAc)^56^ with a correction for spontaneous hydrolysis. The assay mixture (100 µL total volume; path length 0.29 cm) contained 6 mM of *p*NPAc, 8 vol% acetonitrile, 5 mM BES, pH 7.2, and up to 5 µM enzyme. The formation of *p*-nitrophenol/ *p*-nitrophenolate was monitored by the increase in absorbance at 404 nm. Concentrations were calculated using ε_404_ = 11.4 × 10^3^ M^-1^cm^-1^, which accounts for incomplete ionization of *p*-nitro-phenol at pH 7.2. For steady state kinetic measurement, the same assay conditions were used except that the *p*NPAc concentration varied from 0.5 to 8 mM. Kinetic parameters were obtained by fitting the Michaelis−Menten equation by nonlinear regression using either the Solver function of Microsoft Excel or scripts written for the program R^57^. The reported errors are standard deviations. Esterase activity with substrates lacking a chromogenic leaving group were measured using a pH-indicator assay using *p*-nitrophenol as the pH indicator.^58^

The effect of solvent viscosity on steady-state kinetics of *p*-nitrophenyl acetate hydrolysis was determined using the same conditions, but with sucrose as the viscogen at concentrations of 0, 20, 28, and 33 w/w%. Solution viscosities were taken from measured viscosities.^59^ The steady state kinetics used duplicate measurements of approximately ten substrate concentrations of up to 8 mM. The Michaelis-Menten equation was fit to all twenty data points using a non-linear fit to derive the *k*_cat_ and *k*_cat_/*K*_M_ values. The errors indicated are the standard deviations of these values. For HNL1_1st_8, *k*_cat_ did not vary with solvent viscosity and the four data points were fit to a line (slope = 0.00). The other data in Figure 11 showed an increase in *k*_cat_ or in *k*_cat_/*K*_M_ with increasing solvent viscosity, a decrease in ratio. This behavior is not expected to be linear and was not fit to a theoretical line.

Hydroxynitrile lyase activity was measured using mandelonitrile as described previously^21^. The detection limit for *k*_cat_ was 0.1 min^-1^.

### Identification of residues associated with esterase activity

Residues associated with enhanced or decreased esterase activity toward methyl pentanoate at 2 mM substrate concentration were identified by activity-weighted sequence analysis using SigniSite^25^. The sequence alignment and activity values used are in Table S2. The options used for SigniSite were: Exclude gaps from the evaluation = checked (default is unchecked), Significance threshold = 0.05 (default value), Method for correction for multiple testing = no correction (default is Bonferroni single step), Choose sorting of numerical values = decreasing (default), Unique sequence lD for relative numbering = SABP2, Type of logo = Significant residues (default is significant positions), and Include all positions in logo (unchecked, default). The resulting output is shown in Figure 3.

### Homolog search of SABP2

Homologs of SABP2 were identified using the Consensus Finder^60^ web tool (kazlab.umn.edu). An initial blastp search^61^ of the NCBI non-redundant database identified homologs, which were filtered with CD-HIT^62^ at a redundancy setting of 0.90. This filtering clustered the sequences into groups having 90% sequence identity and retained one sequence from each group to yield 864 homologs. This filtering avoids bias from sequences that may be overrepresented in the database. Clustal Omega^63^ created the multiple sequence alignment with 52±21% (mean and standard deviation) identical residues to SABP2. Logo plots of amino acid frequencies were generated from this alignment using Python 3.12 with the logomaker library (v0.8.7) and Biopython (v1.86), excluding gaps from the analysis. The 136 positions identified as identical between SABP2 (Q6RYA0) and HNL1 through pairwise sequence alignment were excluded from logo visualization while maintaining their positions in the plot layout. The resulting logo plot displays amino acid probabilities using a chemistry-based color scheme, with letter height proportional to conservation probability at each position.

### Structural alignment

The Cɑ atoms in the A chains of proteins were aligned using the align function in PyMOL 3. The align function omits unresolved residue pairs and repeats the alignment up to five times and removes outliers at each cycle. Alignment of the structure of HNL1 (PDB ID 5TDX, 260 resolved residues) onto the structure of SABP2 (PDB ID 1Y7H, 257 resolved residues) yielded a root-mean-square deviation of 0.60 Å over 207 atoms, meaning that 50 amino acid residue pairs were removed as outliers. Alignment of the structure of HNL1-15 (PDB ID 9DK4, 248 resolved residues) onto the same SABP2 structure yielded a root-mean-square deviation of 0.55 Å over 208 atoms, meaning that 40 amino acid residue pairs were removed as outliers.

### Docking

Docking was performed with the SwissDock web tool using the Attracting Cavities (AC/AC 2.0) engine^26^. Docking was restricted to a 15-Å cube centered at the Oγ atom of the catalytic serine. The Attracting Cavities approach first replaces the rough interaction surface of the protein by a smooth attractive potential that guides the ligand toward the binding pockets. Ligand poses are then refined on the actual receptor energy landscape, and the resulting complexes are scored using a function based on the CHARMM force field with FACTS solvation. The protein structures are automatically protonated to the form expected at physiological pH before docking. Final poses are clustered into ten clusters, which are ranked by the AC docking score; an additional SwissParam score estimated the binding free energy. Productive orientations were defined as those where the *p*-nitrophenyl moiety of *p*NPAc orients deeper into the cavity than the acetyl moiety; non-productive orientations had the reversed orientation where the acetyl moiety was deeper. Ambiguous orientations placed the *p*NPAc sideways in the pocket.

### Active site cavity volume

Active site cavity volumes were calculated using a grid-based exclusion method implemented in Python 3, following the approach of POVME^64^. The structure HNL1-15 (9DK4, chain A) was superposed onto the structure of HNL1 (5TDX, chain A) by least-squares fitting of 248 shared Cα atoms (RMSD 0.57 Å) using BioPython 1.86. The active site was defined by a spherical inclusion region of radius 7.0 Å centered on the centroid of the glycerol molecule (residue GOL 300) in the 5DTX structure. A uniform grid of points at 1.0 Å spacing was generated within this sphere. Grid points were excluded if they fell within the van der Waals radius of any protein heavy atom (C 1.70, N 1.55, O 1.52, S 1.80 Å), or if no protein heavy atom was present within 4.0 Å (to exclude solvent-exposed exterior points not enclosed by the protein). Cavity volume was computed as the number of remaining grid points multiplied by the grid spacing cubed (1.0 Å³ per point). To identify cavity volume gained or lost between the two structures, grid points unique to each cavity were identified using a distance threshold of 0.75 Å via scipy cKDTree nearest-neighbor search.

### Tunnels

Tunnels were calculated using CAVER 3.0^17^ command-line version starting from pdb structures (1Y7I for SABP2, 5TDX for HNL1, and 9DK4 for HNL1-15). The structures were further prepared by removing water, non-protein HETATM such as salicylate for SABP2 and glycerol for HNL1, and all protein chains except chain A for SABP2 and HNL1. For HNL1-15 we used chain B because it had fewer missing residues. The structure of SABP2 contained nine selenomethionine residues, which were converted to methionine by renaming the residue and relabeling the selenium atom as sulfur. The structure of HNL1 contained alternative positions for 78 atoms, which were removed leaving only the major positions. The structure of HNL1-15, chain B, was missing residues 70-72, so they were modeled with SwissModel^65^ using the 9DK4 structure as the template. The tunnel-search starting point was the Oγ of the catalytic serine. A minimum probe radius of 0.86 Å was used with a shell radius of 3 Å and shell depth of 4. Tunnels with a cost above 1.4 were discarded as likely spurious, leaving three tunnels for SABP2, one for HNL1 and two for HNL1-15.

### B-factor normalization

To compare backbone flexibility in HNL1 and HNL1-15, Cα B-factors were converted to Z-scores within each structure by subtracting the mean and dividing by the standard deviation over the 248 residues common to both models. Per-residue differences were then calculated as ΔZ = Z_9DK4_ − Z_5TDX_. Because the two structures were refined independently and differ in resolution and crystal packing, raw B-factors are not directly comparable. This normalization places both structures on a common scale of relative mobility, so that ΔZ reports whether a residue is more or less mobile than average within *its own structure*. The sets of the twenty largest increases and decreases in ΔZ identified the regions shown in Figure S3.

## Supporting Information

Additional data (23 pages) consisting of four figures, ten tables and nine amino acid sequences.

## Acknowledgments

The authors thank Prof. Silvia Osuna (ICREA, Universitat de Girona) and Dr. Guillem Casadevall (Universitat de Girona) for helpful discussions. This research was supported by the National Institutes of Health/National Institute of General Medical Sciences (grant No. GM119483 and R35-GM118047) and by the National Science Foundation (NSF Award No. CBET-2039039). This research used the beam line of the National Synchrotron Light Source II, a U.S. Department of Energy (DOE) Office of Science User Facility operated for the DOE Office of Science by Brookhaven National Laboratory under Contract No. DE-SC0012704. The Center for BioMolecular Structure (CBMS) is primarily supported by the National Institutes of Health, National Institute of General Medical Sciences (NIGMS) through a Center Core P30 Grant (P30GM133893), and by the DOE Office of Biological and Environmental Research (KP1605010). The contents of this publication are solely the responsibility of the authors and do not necessarily represent the official views of NIGMS or NIH.

