## Supplemental Information for "Identifying a cooperative catalytic network for efficient esterase catalysis"

**Contents**

**Figure S1**. Mechanism for hydrolysis of *p*-nitrophenyl acetate catalyzed by a serine esterase.
**Figure S2.** Sequence alignment of selected enzymes.
**Figure S3.** Normalized flexibility changes in HNL1-15 as compared to HNL1.
**Figure S4.** Sequence logo plot of an MSAof 864 homologs of SABP2

**Table S1.** Steady-state kinetic constants for HNL1 variants and selected other enzymes.
**Table S2.** Sequence alignment of the ten sequences used in the SigniSite analysis.
**Table S3.** Thirty-eight substitutions in HNL1 predicted to increase activity toward pNPAc.
**Table S4.** Steady-state kinetic constants for HNL1 variants toward differently shaped esters.
**Table S5.** Catalytic activity toward *p*NPAc of HNL1-16 reversion variants.
**Table S6.** HNL1-15 protein production information.
**Table S7.** Crystallization information for HNL1-15.
**Table S8.** Data collection and processing for HNL1-15.
**Table S9.** Structure refinement for HNL1-15 (pdb id 9DK4).
**Table S10.** Calculated tunnel characteristics in SABP2, HNL1 and HNL1-15.

**Amino acid sequences**

### Figures

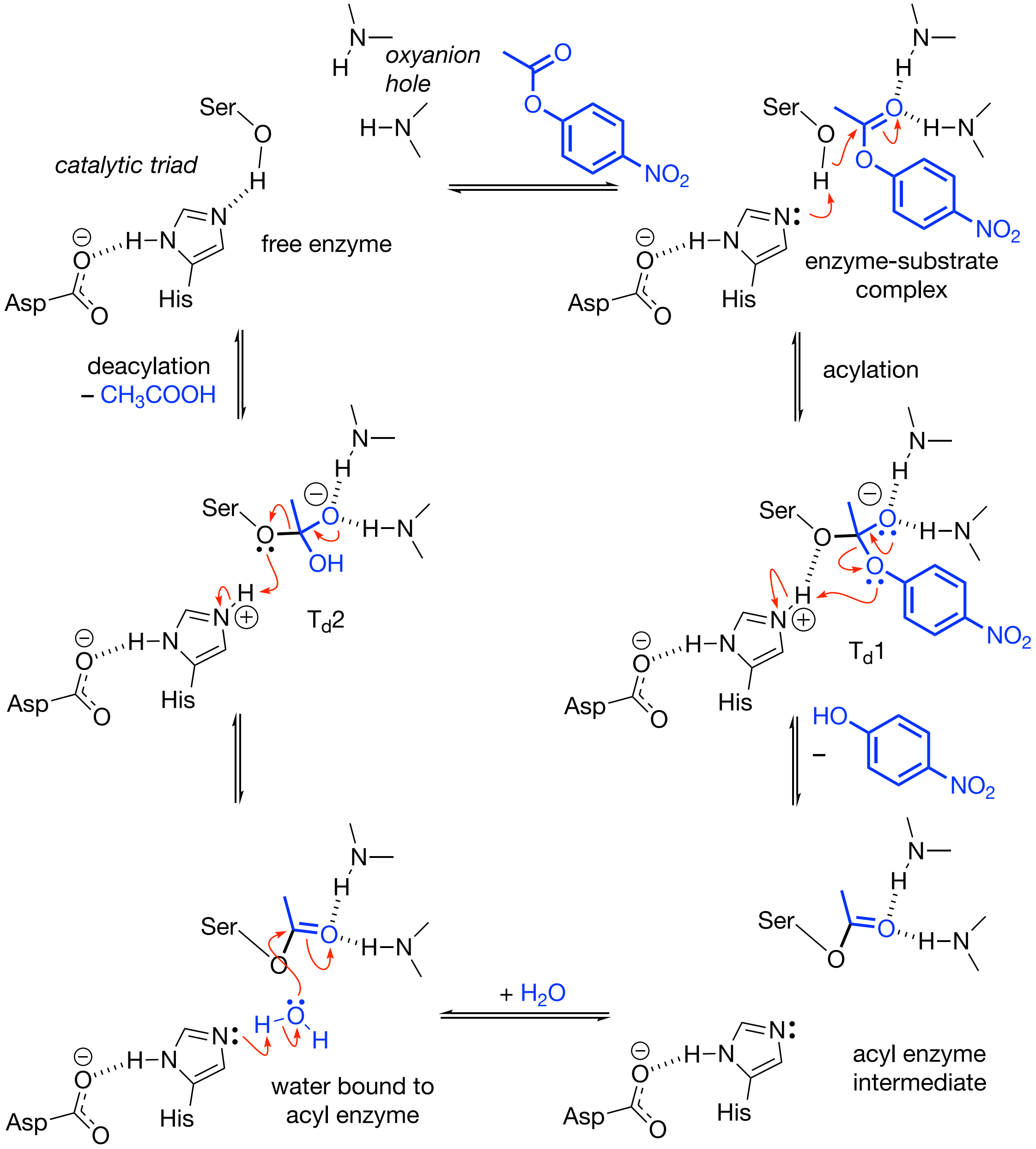


**Figure S1.** Reaction mechanism for hydrolysis of *p*-nitrophenyl acetate catalyzed by a serine esterase.

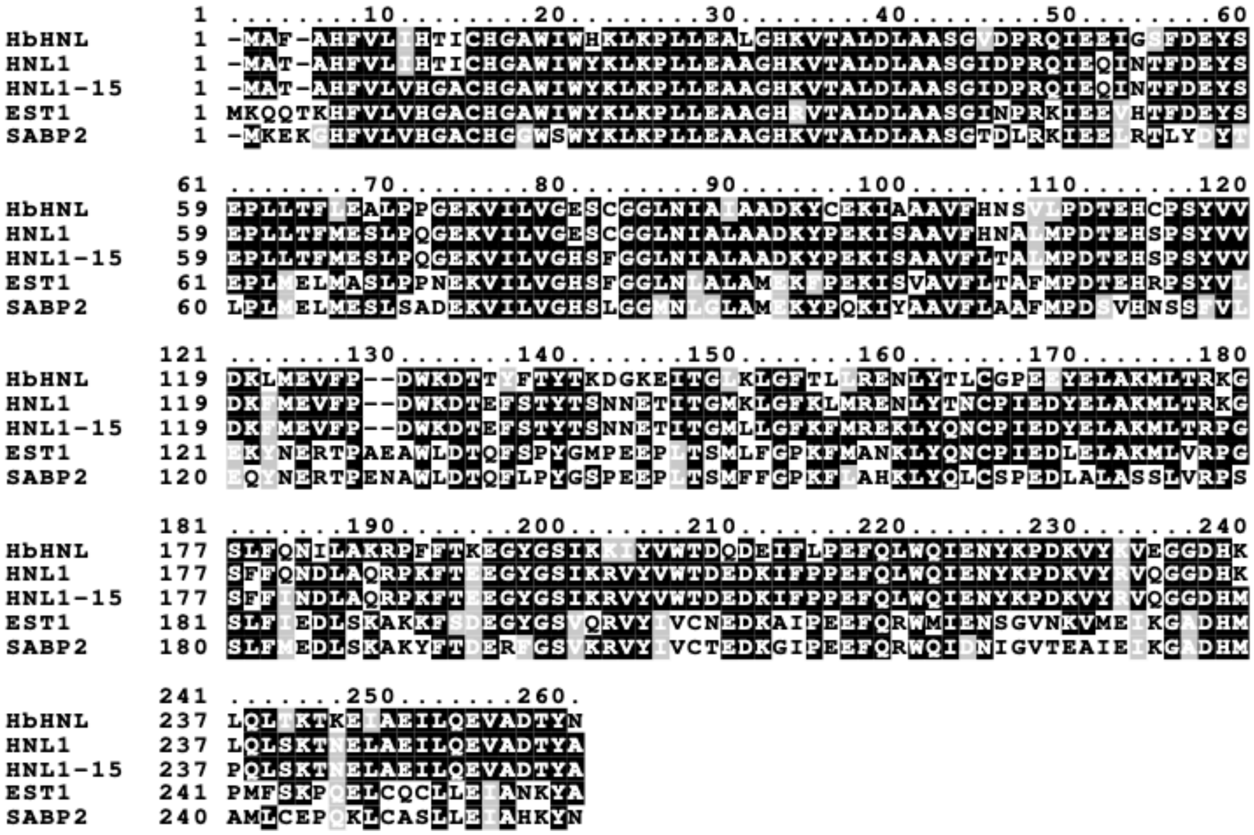


**Figure S2.** Sequence alignment of selected proteins. Positions with identical amino acids are shaded black, positions with similar amino acids are shaded gray and positions with dissimilar amino acids are unshaded. The catalytic triad is conserved in all five proteins and occurs at Ser80–His235–Asp207 in the HNL numbering. Sequences were aligned with Clustal Omega (Sievers et al., 2011) and shaded with Boxshade (https://junli.netlify.app/apps/boxshade/). The proteins contained a C-terminal His6-tag which is not shown in this alignment

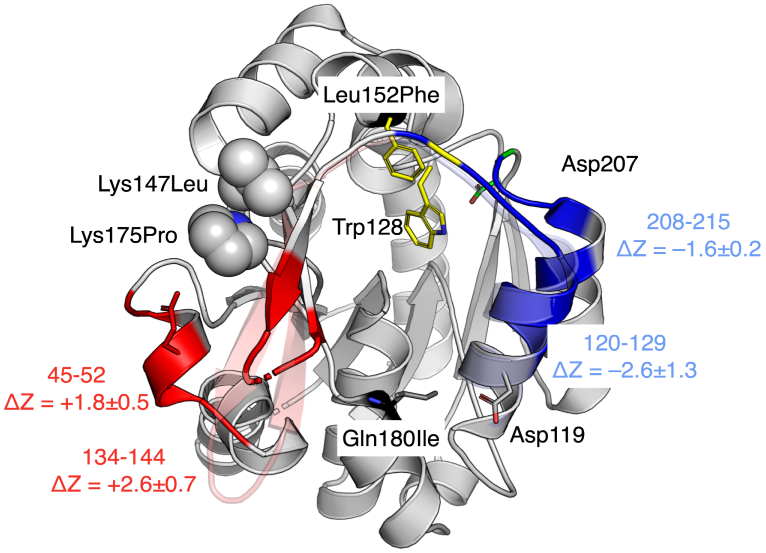


**Figure S3.** Normalized flexibility changes in HNL1-15 as compared to HNL1. HNL1-15 (9DK4, solid cartoon) is superimposed on HNL1 (5TDX). Regions of increased flexibility in HNL1-15 are colored red (loop-helix-loop 45–52 and strand-loop-strand 134–144); regions of decreased flexibility are colored blue (helix-loop 120–129 and helix-loop 208–215). The ghost cartoons show the corresponding regions of HNL1: the light blue ghost illustrates the positional shift of helix 115–132, and the salmon ghost indicates the 134–144 loop that is ordered in HNL1 but partially disordered in HNL1-15. Black spheres mark the Cα positions of substitution sites discussed in the text. Selected sidechains are shown as sticks. The rigidified helix-loop 208-215 may fix the position of the catalytic triad residue Asp207 (green sticks).

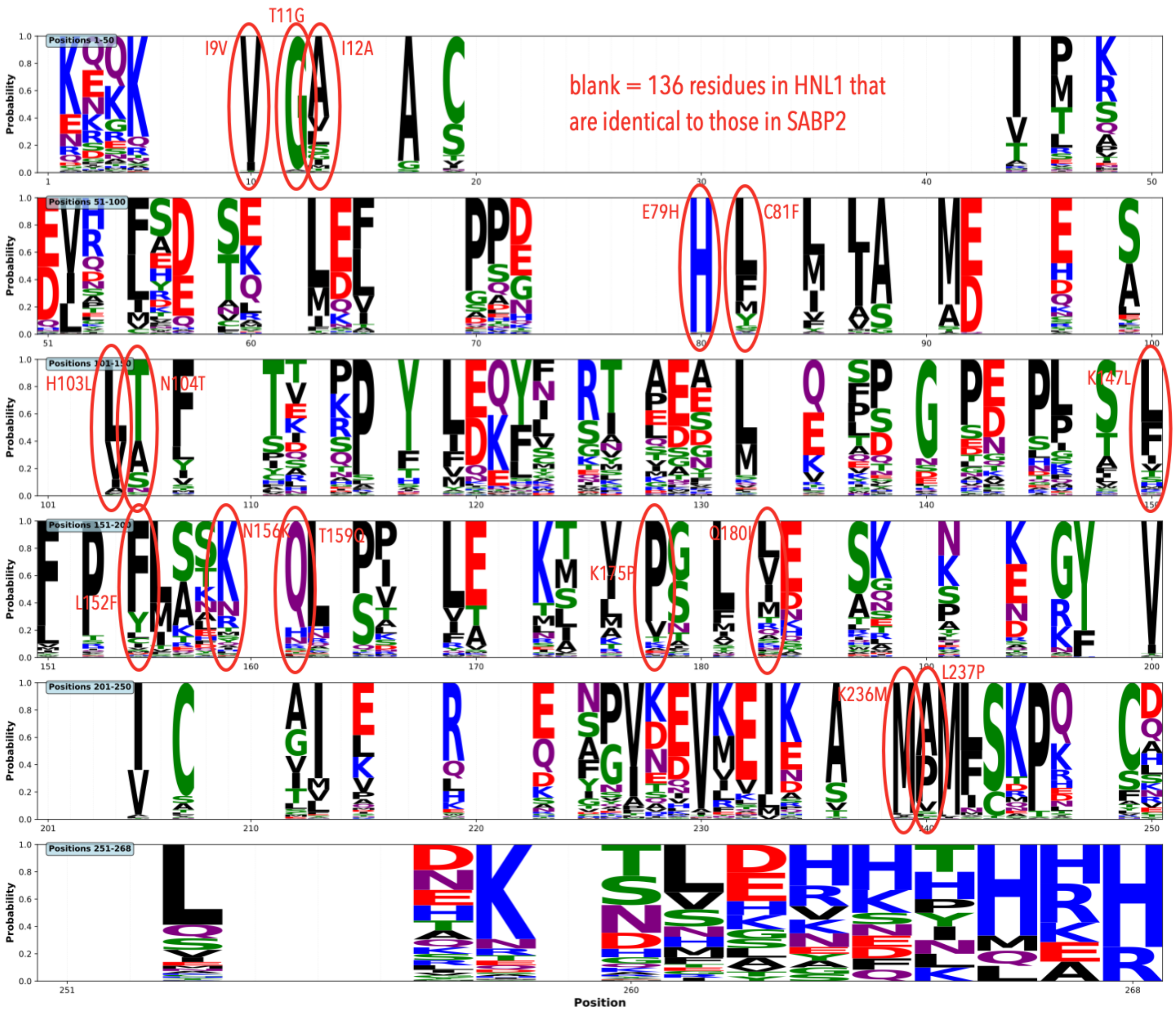


**Figure S4.** Sequence logo plot of a multiple sequence alignment of 864 homologs of SABP2 showing that eleven of the fifteen catalytically essential substitutions are not conserved in SABP2 homologs. The average conservation over all 260 SABP2 positions among the 864 homologs is 52±21%. The x-axis numbers the amino acids as in SABP2; the labels use HNL1 numbering. The residues identical in both HNL1 and SABP2 (136 positions, 51.5%) are not shown for clarity. For the remaining 124 positions, the letter height indicates the probability of each amino acid at that position and the colors represent amino acid chemical properties. The red ovals mark the 15 amino acids that were substituted in HNL1 to create HNL1-15. At four positions (I9V, T11G, E79H and K236M, HNL1 numbering) the replacement amino acid occurs in >90% of the homologs, but at the other eleven positions, the replacement amino acid conservation matches the average conservation over all positions.

**Tables**

**Table S1.** Steady-state kinetic constants for HNL1 variants and selected other enzymes of the hydrolysis of *p*-nitrophenyl acetate. These values are presented graphically in Figure 2 of the main text.

| **Enzyme name** | ***p*-Nitrophenyl acetate** | | |
|---|---|---|---|
| | $k_{cat}$ (min$^{-1}$) | $K_M$ (mM) | $k_{cat}/K_M$ (min$^{-1}$ M$^{-1}$) |
| *Hb*HNL | 0.3±0.02 | 3.0±0.4 | 100 |
| SABP2 | 134±4 | 2.2±0.2 | 61,000 |
| HNL1 | 1.7±0.1 | 9±1 | 190 |
| EST1 | 140±2.5 | 1.8±0.1 | 77,000 |
| HNL1_SigniSite_38 | 23±0.6 | 0.5±0.1 | 49,000 |
| HNL1_all_close_15 | 1.6±0.1 | 1.7±0.3 | 920 |
| HNL1_1st_8 | 7.9±0.5 | 1.2±2 | 6,400 |
| HNL1_2nd_8 | *a* | *a* | 97 |
| HNL 1-16 | 73±1.5 | 0.8±0.1 | 87,000 |
| HNL1-15 | 109±2.1 | 1.3±0.1 | 81,000 |

[a]The rate of HNL1_2nd_8res-catalyzed hydrolysis of *p*NPAc varied linearly with substrate concentration up to the solubility limit of 8 mM. The slope of this line was 97 min$^{-1}$ M$^{-1}$ and corresponds to the ratio of $k_{cat}/K_M$. The steady state kinetic constants $k_{cat}$ and $K_M$ could not be determined individually since the enzyme showed no evidence of saturation at the highest substrate concentrations tested.

**Table S2.** Sequence alignment of the ten sequences used in the SigniSite analysis to identify residues associated with increased or decreased esterase activity. The numbers after the protein name are the measured esterase activity ($min^{-1}$) at 2 mM methyl pentanoate from Table S2 in Devamani *et al.*[18]. Esterase activity was monitored spectrophotometrically using *p*-nitrophenol as the pH indicator.

```
>EST2 340
------MAEMKNRTKHFVLVHGACHGAWVWYKLKPLLEAAGHRVTALDLAASGINPKKIE
EVHTFDEYSEPLMELMASLPPNEKVILVGHSLGGLNLALAMEKFPEKISVAVFLTAFMPD
TEHRPSYVLEKYNERTPAEAWLDTQFSPYGNPEEP-LTSMLFGPKFMANKLYQLSPIEDL
ELAKMLVRPGSLF-IEDLSKAKKFSDEGYGSVPRVYIVCNEDKAIPEEFQRWMIENSGVN
EVMEIKGADHMPMFSKPQELCQCLLEIANKYAKAGDPLGGG
>SABP2 198
----------KEG-KHFVLVHGACHGGWSWYKLKPLLEAAGHKVTALDLAASGTDLRKIE
ELRTLYDYTLPLMELMESLSADEKVILVGHSLGGMNLGLAMEKYPQKIYAAVFLAAFMPD
SVHNSSFVLEQYNERTPAENWLDTQFLPYGSPEEP-LTSMFFGPKFLAHKLYQLCSPEDL
ALASSLVRPSSLF-MEDLSKAKYFTDERFGSVKRVYIVCTEDKGIPEEFQRWQIDNIGVT
EAIEIKGADHMAMLCEPQKLCASLLEIAHKYN---------
>EST1 140
------MAEMKQQTKHFVLVHGACHGAWIWYKLKPLLEAAGHRVTALDLAASGINPRKIE
EVHTFDEYSEPLMELMASLPPNEKVILVGHSFGGLNLALAMEKFPEKISVAVFLTAFMPD
TEHRPSYVLEKYNERTPAEAWLDTQFSPYGMPEEP-LTSMLFGPKFMANKLYQNCPIEDL
ELAKMLVRPGSLF-IEDLSKAKKFSDEGYGSVQRVYIVCNEDKAIPEEFQRWMIENSGVN
KVMEIKGADHMPMFSKPQELCQCLLEIANKYAKAGDPLGGG
>RsEST 7.1
----MHSAANAKQQKHFVLVHGGCLGAWIWYKLKPLLESAGHKVTAVDLSAAGINPRRLD
EIHTFRDYSEPLMEVMASIPPDEKVVLLGHSFGGMSLGLAMETYPEKISVAVFMSAMMPD
PNHSLTYPFEKYNEKCPADMMLDSQFSTYGNPENP-GMSMILGPQFMALKMFQNCSVEDL
ELAKMLTRPGSLF-FQDLAKAKKFSTERYGSVKRAYIFCNEDKSFPVEFQKWFVESVGAD
KVKEIKEADHMGMLSQPREVCKCLLDISDS-----------
>RcEST 2.8
-GKQVPDFAENIKFKKFILVHGEGFGAWCWYKTVALLEEAGLLPTALDLTGSGIHLTDTN
SVTKLADYSQPLINYLENLPEDEKVILVGHSTGGACISLALEHFPQKISKAIFLCATMVS
DGQRPFDVFAEELGSA-ERFMQESEFLIYGNGKDKAPTGFMFEKQQMKGLYFNQSTTKDV
ALAMVCMRPIPLG---PVMEKLSLSPEKYGTGRRFFIQTLDDHALSPDVQEKLVRENPPE
GVFKIKGSDHCPFFSKPQSLHKILLEIAQIP----------
>HNL1-NJ 1
-----------MAVAHFVLIHTICHGAWIWYKLKPLLESAGHKVTALDLAASGIDPRQIE
QVGTFEEYSEPLLTFLESLPEGEKVILVGESCGGINIALAADKYPEKISAAVFHNALMPD
TVHSPSYVLDKMFEVFP--DWKDSVFSNYTNGSNDTITALKLGPKLMKENIYTNCPIEDY
ELAKMLVRKGSLF-QEDLAKREKFTEEGYGSIKRVYVYGDEDKIFLEEFQRWQINNYKPD
KVYEVPGGDHKLMLSKVNELFQILQEVADTYASLLAVAGGG
>HNL1-ML 0.45
-----------MAWAHFVLIHTICHGAWIWYKLKPLLEAAGHKVTALDLAASGIDPRQIE
QIGSFDEYSEPLLTFMESLPQGEKVILVGESCGGINIAIAADKYPEKIAAAVFHNALMPD
```

TVHNPSYVLDKFMEVFP--DWKDSEFSNYTYG-NDTITALKLGPKLMKENLYTNCPPEDY
ELAKMLVRKGSLF-QEDLAKRENFTKEGYGSIKRIYVYGDEDKIFTEEFQRWQIDNYKPD
KVYVVPGGDHKLMLSKVNELFQILQEVADTYANLLAVGGGH
>HbHNL 0.44
-----------MAFAHFVLIHTICHGAWIWHKLKPLLEALGHKVTALDLAASGVDPRQIE
EIGSFDEYSEPLLTFLEALPPGEKVILVGESCGGLNIAIAADKYCEKIAAAVFHNSVLPD
TEHCPSYVVDKLMEVFP--DWKDTTYFTYTKD-GKEITGLKLGFTLLRENLYTLCGPEEY
ELAKMLTRKGSLF-QNILAKRPFFTKEGYGSIKKIYVWTDQDEIFLPEFQLWQIENYKPD
KVYKVEGGDHKLQLTKTKEIAEILQEVADTYN---------
>MeHNL 0.44
-----------MVTAHFVLIHTICHGAWIWHKLKPALERAGHKVTALDMAASGIDPRQIE
QINSFDEYSEPLLTFLEKLPQGEKVIIVGESCAGLNIAIAADRYVDKIAAGVFHNSLLPD
TVHSPSYTVEKLLESLP--DWRDTEYFTFTNITGETITTMKLGFVLLRENLFTKCTDGEY
ELAKMVMRKGSLF-QNVLAQRPKFTEKGYGSIKKVYIWTDQDKVFLPDFQRWQIANYKPD
KAYQVQGGDHKLQLTKTEEVAHILQEVADAYA---------
>HNL1 0.44
-----------MATAHFVLIHTICHGAWIWYKLKPLLEAAGHKVTALDLAASGIDPRQIE
QINTFDEYSEPLLTFMESLPQGEKVILVGESCGGLNIALAADKYPEKISAAVFHNALMPD
TEHSPSYVVDKFMEVFP--DWKDTEFSTYTSN-NETITGMKLGFKLMRENLYTNCPIEDY
ELAKMLTRKGSFF-QNDLAQRPKFTEEGYGSIKRVYVWTDEDKIFPPEFQLWQIENYKPD
KVYRVQGGDHKLQLSKTNELAEILQEVADTYADLLAVAGGG

**Table S3.** Thirty-eight substitutions in HNL1 predicted to increase esterase activity. The locations are shown in Figure 4 of the main text.

| | Substitution in HNL1 | location relative to active site |
|---|---|---|
| 1 | A4K | 3rd shell (> 10.5 Å) |
| 2 | I9V | 2nd shell (7-10.5 Å) |
| 3 | T11G | 1st shell (<7 Å) |
| 4 | I12A | 1st shell (<7 Å) |
| 5 | Q47K | 3rd shell (> 10.5 Å) |
| 6 | L62M | 3rd shell (> 10.5 Å) |
| 7 | T63E | 3rd shell (> 10.5 Å) |
| 8 | F64L | 3rd shell (> 10.5 Å) |
| 9 | G71N | 3rd shell (> 10.5 Å) |
| 10 | E79H | 1st shell (<7 Å) |
| 11 | C81F | 1st shell (<7 Å) |
| 12 | A90M | 3rd shell (> 10.5 Å) |
| 13 | D91E | 3rd shell (> 10.5 Å) |

| | | |
|---|---|---|
| 14 | H103L | 2nd shell (7-10.5 Å) |
| 15 | N104T | 1st shell (<7 Å) |
| 16 | T137G | 3rd shell (> 10.5 Å) |
| 17 | I143L | 3rd shell (> 10.5 Å) |
| 18 | K147L | 2nd shell (7-10.5 Å) |
| 19 | L152F | 1st shell (<7 Å) |
| 20 | E155N | 2nd shell (7-10.5 Å) |
| 21 | N156K | 1st shell (<7 Å) |
| 22 | T159Q | 2nd shell (7-10.5 Å) |
| 23 | Y166L | 3rd shell (> 10.5 Å) |
| 24 | K175P | 2nd shell (7-10.5 Å) |
| 25 | Q180I | 2nd shell (7-10.5 Å) |
| 26 | R186A | 3rd shell (> 10.5 Å) |
| 27 | I197V | 3rd shell (> 10.5 Å) |
| 28 | D205N | 3rd shell (> 10.5 Å) |
| 29 | Y222S | 3rd shell (> 10.5 Å) |

| | | |
|---|---|---|
| 30 | K223G | 3rd shell (> 10.5 Å) |
| 31 | Y228M | 3rd shell (> 10.5 Å) |
| 32 | V230I | 3rd shell (> 10.5 Å) |
| 33 | G233A | 3rd shell (> 10.5 Å) |
| 34 | K236M | 1st shell (<7 Å) |
| 35 | L237P | 2nd shell (7-10.5 Å) |
| 36 | T242P | 3rd shell (> 10.5 Å) |
| 37 | Q250L | 3rd shell (> 10.5 Å) |
| 38 | V252I | 3rd shell (> 10.5 Å) |

**Table S4.** Steady-state kinetic parameters for hydrolysis of esters of similar overall size, but different shapes.

| Substrate | Enzyme | $K_M$ (mM) | $k_{cat}$ ($min^{-1}$) | relative $k_{cat}$ |
|---|---|---|---|---|
| *p*-nitrophenyl acetate (largest alcohol) | HNL1 | 8.7±1.1 | 1.7±0.1 | 1 |
| | HNL1-16 | 0.8±0.1 | 73±2 | 43 |
| phenyl formate (large alcohol) | HNL1 | 0.24±0.08 | 5.9±0.3 | 1 |
| | HNL1-16 | 0.34±0.06 | 46±1 | 7.7 |
| methyl phenylacetate (small alcohol) | HNL1 | 0.09±0.05 | 8.7±0.6 | 1 |
| | HNL1-16 | 0.4±0.1 | 18±2 | 2.0 |
| methyl benzoate (small alcohol) | HNL1 | 0.5±0.2 | 3.7±0.3 | 1 |
| | HNL1-16 | 0.14±0.05 | 12±1 | 3.2 |

**Table S5.** Catalytic activity toward *p*NPAc of HNL1-16 reversion variants where one of the substitutions is restored to the original amino acid residue in HNL1. The standard errors are from a best fit to the Michaelis-Menten equation of three replicate measurements. This data is also shown in Figure 5 of the main text.

| **Mutation** | $k_{cat}$ (min$^{-1}$) | $K_M$ (mM) | $k_{cat}/K_M$ (min$^{-1}$ M$^{-1}$) | $k_{cat}$ relative to HNL1-16 |
|---|---|---|---|---|
| HNL1-16 | 72.6±1.5 | 0.8±0.1 | 87000 | 100% |
| **1st Shell (<7.0 Å)** | | | | |
| A12I | 2.8±0.1 | 0.5±0.02 | 5100 | 4% |
| F81C | 0.1±0.1 | 0.5±0.3 | 200 | 0% |
| M236K | 0.01±0 | 0.3±0.1 | 33 | 0% |
| F152L | 28±0.5 | 0.2±0.02 | 140000 | 39% |
| G11T | 0.01±0 | 2.8±0.5 | 4 | 0% |
| H79E | 0.01±0 | 2.0±0.2 | 5 | 0% |
| K156N | 6.9±0.2 | 2.5±0.1 | 2760 | 9% |
| T104N | 2.0±0.1 | 3.2±0.3 | 625 | 3% |
| **2nd Shell (7–10.5 Å)** | | | | |
| L147K | 0.16±0 | 1.8±0.8 | 89 | 0% |
| N155E = HNL1-15 | 109±2.1 | 1.3±0.1 | 81000 | 150% |
| P175K | 1.04±0.1 | 0.8±0.1 | 1200 | 1% |
| P237L | 3.3±0.1 | 0.8±0.1 | 4100 | 5% |
| I180Q | 2.4±0 | 0.8±0.02 | 3000 | 3% |
| Q159T | 12.3±0.3 | 1.1±0.1 | 11100 | 17% |
| L103H | 0.21±0 | 0.9±0.1 | 250 | 0% |
| V9I | 5.0±0.1 | 1.0±0.1 | 5000 | 7% |
| **Remove two substitutions** | | | | |

| Mutation | $k_{cat}$ ($min^{-1}$) | $K_M$ (mM) | $k_{cat}/K_M$ ($min^{-1}$ $M^{-1}$) | $k_{cat}$ relative to HNL1-16 |
|---|---|---|---|---|
| HNL1-14 (L103H, T104N) | 5.6±0.1 | 0.4±0.03 | 14000 | 8% |

**Table S6.** HNL1-15 protein production information.

| | |
|---|---|
| Source Organism | Synthetic construct |
| DNA source | Synthetic construct |
| Expression vector | pET21a(+) |
| Expression host | *E. coli* BL21 |
| Complete amino-acid sequence of the construct produced | MATAHFVLVHGACHGAWIWYKLKPL-LEAAGHKVTALDLAASGID-PRQIEQINTFDEYSE-PLLTFMESLPQGEKVILVGHSFGGL-NIALAADKYPEKISAAVFLTALM-PDTEHSPSYVVDKFMEVFPDWKDTEF-STYTSNNETITGMLLGFKFMREK-LYQNCPIEDYELAKMLTRPGSFFIND-LAQRPKFTEEGYGSIKRVYVWTDED-KIFPPEFQLWQIENYKPDKVYRVQG-GDHMPQLSKTNELAEILQE-VADTYADLLAVAGLEHHHHHH |

| | |
|---|---|
| Codon-optimized coding DNA sequence (CDS) | ATGGCAACCGCTCATTTTGTATTAGTA-<br>CACGGTGCATGCCACGGCGCATG-<br>GATTTGGTATAAGCTCAAGCCACTTT-<br>TAGAAGCTGCCGGACATAAAGTAACT-<br>GCCTTAGATTTGGCAGCGAGCG-<br>GTATAGATCCGCGTCAGATAGAACA-<br>GATCAATACCTTCGATGAATACTCA-<br>GAGCCACTTCTGACCTTTATG-<br>GAATCGCTACCCCAAGGAGAAAAG-<br>GTAATACTTGTTGGTCATTCTTTTG-<br>GAGGACTGAACATCGCATTGGCTGCA-<br>GATAAATATCCCGAAAAAATTAGT-<br>GCAGCGGTGTTTCTTACAGCCCTTAT-<br>GCCGGATACAGAACACAGTCCGTCC-<br>TATGTGGTAGATAAATTTATGGAG-<br>GTCTTTCCGGATTGGAAAGATACT-<br>GAATTCTCCACGTATACTAGCAATAAC-<br>GAGACCATCACCGGCATGCTTCTTG-<br>GCTTTAAATTTATGCGGAAAAATTG-<br>TACCAAAACTGCCCAATCGAGGAT-<br>TATGAGCTGGCAAAAATGCT-<br>GACGCGTCCGGGGAGTTTTTTAT-<br>TAACGATTTGGCACAACGTCCTAAGTT-<br>TACCGAAGAGGGTTACGGGTCTAT-<br>TAAAAGAGTATATGTATGGACAGAC-<br>GAAGACAAAATTTTTCCGCCA-<br>GAATTTCAATTATGGCAAATAGAGAAC-<br>TACAAACCGGATAAAGTC-<br>TACAGGGTACAGGGTGGTGACCATAT-<br>GCCGCAGCTGTCTAAAACCAAC-<br>GAATTGGCAGAAATTCT-<br>GCAGGAAGTGGCAGATACCTACGCG-<br>GATCTGCTTGCTGTGGCCGGTCTC-<br>GAGCACCACCACCACCACCACTGA |

**Table S7.** Crystallization information for HNL1-15.

| | |
|---|---|
| Method | Sitting Drop |
| Plate type | CrystalMation Intelli-Plate low-profile 96 Well, Hampton Research |
| Temperature (K) | 298 |
| Protein concentration (mg $mL^{-1}$) | 5.0 |
| Buffer composition of protein solution | 5 mM BES buffer, pH 7.2 |
| Composition of reservoir solution | 5%(w/v) PEG 20K, 25% (w/v) 1,1,1-tris(hydroxymethyl)propane, 1% (w/v) NDSB 195 (dimethylethylammonium propane sulfonate), 1 mM of each alkali (barium acetate, cesium acetate, rubidium chloride, strontium acetate) and 0.1 M *N*,*N*-bis(2-hydroxyethyl)-2-aminoethanesulfonic acid/triethylamine pH 7.5 |
| Volume and ratio of drop (μL) | 0.1, 1:1 |
| Volume of reservoir (μL) | 50 |

**Table S8.** Data collection and processing for HNL1-15.

Values in parentheses are for the highest-resolution shell

| | |
|---|---|
| X-ray source | NSLSII-17-ID-2 |
| Wavelength (Å) | 0.979 |
| Detector | Eiger16M |
| Exposure Time (s) | 0.05 |
| Crystal-to-detector distance (mm) | 300 |
| Angle increment (°) | 0.2 |
| Resolution Range (Å) | 46.41 - 1.88 (1.947 - 1.88) |
| Space Group | $P2_1$ |
| a, b, c (Å) | 67.244 80.385 92.823 |
| α, β, γ (°) | 90 90.622 90 |
| Matthews coefficient ($Å^3Da^{-1}$) | 2.03 |
| Solvent Content (%) | 39.36 |
| Total reflections | 234102 (10618) |
| Unique Reflections | 73427 (4798) |
| Multiplicity | 3.2 (2.9) |
| Mosaicity (°) | 0.2 |
| Completeness (%) | 91.22 (59.86) |
| ($I/\sigma(I)$) | 4.6 (1.9) |

| | |
|---|---|
| Wilson B Factor (Å$^2$) | 18.59 |
| $R_{merge}$ | 0.137 (0.408) |
| $R_{meas}$ | 0.163 (0.493) |
| $R_{p.i.m}$ | 0.087 (0.273) |
| $CC_{1/2}$ | 0.967 (0.715) |

**Table S9.** Structure refinement information for HNL1-15 (pdb id 9DK4).

Values in parentheses are for the highest-resolution shell

| | |
|---|---|
| Reflections used in refinement | 73413 (4794) |
| Reflections used for $R_{free}$ | 1993 (136) |
| $R_{work}$ | 0.1837 (0.2564) |
| $R_{free}$ | 0.2283 (0.3178) |
| No. of non-H atoms | |
| Total | 8143 |
| Macromolecules | 7873 |
| Ligands | 7 |
| Solvent | 263 |
| No. of protein residues | 1006 |
| R.m.s.d, bonds (Å) | 0.009 |
| R.m.s.d, angles (°) | 0.94 |
| Ramachandran favored (%) | 96.36 |
| Ramachandran allowed (%) | 3.64 |
| Ramachandran outliers (%) | 0.00 |
| Rotomer outliers (%) | 0.25 |
| Clashscore | 3.08 |
| Average B-factor | 18.88 |
| Macromolecules | 18.80 |
| Ligands | 29.9 |
| Solvent | 20.87 |

**Table S10.** Calculated tunnel characteristics in SABP2, HNL1 and HNL1-15 using Caver. The cost is a geometric accessibility score that takes into account the tunnel width (radius) and length; lower is better; <0.7 is favorable. The throughput is the Caver ranking score and is defined as $e^{-cost}$. Higher throughput is better; > 0.5 is favorable.

| enzyme | name | throughput | cost | radius, Å | length, Å | curvature |
|---|---|---|---|---|---|---|
| SABP2 | right tunnel | 0.62 | 0.47 | 1.2 | 15 | 1.2 |
| SABP2 | left tunnel | 0.60 | 0.51 | 0.92 | 12 | 1.2 |
| SABP2 | back tunnel | 0.33 | 1.1 | 0.92 | 21 | 1.3 |
| HNL1 | left tunnel | 0.52 | 0.65 | 1.2 | 14 | 1.5 |
| HNL1-15 | left tunnel | 0.52 | 0.65 | 1.2 | 17 | 1.7 |
| HNL1-15 | right tunnel | 0.42 | 0.87 | 0.88 | 16 | 1.3 |

**Amino acid sequences**

```
>SABP2 = UniProt Q6RYA0 SABP2_TOBAC
MKEGKHFVLVHGACHGGWSWYKLKPLLEAAGHKVTALDLAASGTDLRKIEELRTLY-
DYTLPLMELMESLSADEKVILVGHSLGGMNLGLAMEKYPQKIYAAVFLAAFMPDSVHNSS-
FVLEQYNERTPAENWLDTQFLPYGSPEEPLTSMFFGPKFLAHKLYQLCSPEDLALASS-
LVRPSSLFMEDLSKAKYFTDERFGSVKRVYIVCTEDKGIPEEFQRWQIDNIGVTEAIEIK-
GADHMAMLCEPQKLCASLLEIAHKYN

>EST1 ancestral esterase
MAEMKQQTKHFVLVHGACHGAWIWYKLKPLLEAAGHRVTALDLAASGINPRKIEEVHTFDEY-
SEPLMELMASLPPNEKVILVGHSFGGLNLALAMEKFPEKISVAVFLTAFMPDTEHRPSYVLE-
KYNERTPAEAWLDTQFSPYGMPEEPLTSMLFGPKFMANKLYQNCPIEDLEL-
AKMLVRPGSLFIEDLSKAKKFSDEGYGSVQRVYIVCNEDKAIPEEFQRWMIENSGVNKVMEIK-
GADHMPMFSKPQELCQCLLEIANKYAKAGDPLGGG

>HbHNL = UniProt P52704 HNL_HEVBR
MAFAHFVLIHTICHGAWIWHKLKPLLEALGHKVTALDLAASGVDPRQIEEIGSFDEYSEPLLT-
FLEALPPGEKVILVGESCGGLNIAIAADKYCEKIAAAVFHNSVLPDTEHCPSYVVDKLMEVF-
PDWKDTTYFTYTKDGKEITGLKLGFTLLRENLYTLCGPEEYELAKMLTRKGSLFQNI-
LAKRPFFTKEGYGSIKKIYVWTDQDEIFLPEFQLWQIENYKPDKVYKVEGGDHKLQLTK-
TKEIAEILQEVADTYN
```

>HNL1 = pdb id 5TDX ancestral hydroxynitrile lyase
MATAHFVLIHTICHGAWIWYKLKPLLEAAGHKVTALDLAASGIDPRQIEQINTFDEYSE-
PLLTFMESLPQGEKVILVGESCGGLNIALAADKYPEKISAAVFHNALMPDTEHSPSYVVD-
KFMEVFPDWKDTEFSTYTSNNETITGMKLGFKLMRENLYTNCPIEDYELAKMLTRKGSFFQND-
LAQRPKFTEEGYGSIKRVYVWTDEDKIFPPEFQLWQIENYKPDKVYRVQGGDHKLQLSKT-
NELAEILQEVADTYADLLAVAGGG

>HNL1-all_close_15 (exchanged all differing aa <7 Å between HNL1 and EST1) T11G, I12A, E79H, C81F, N104T, L106F, F121Y, M122N, L148F, L152F, N156K, F178L, I209A, F210I, K236M
MATAHFVLIHGACHGAWIWYKLKPLLEAAGHKVTALDLAASGIDPRQIEQINTFDEYSEPLL-
TFMESLPQGEKVILVGHSFGGLNIALAADKYPEKISAAVFHTAFMPDTEHSPSYVVDKY-
NEVFPDWKDTEFSTYTSNNETITGMKFGFKFMREKLYTNCPIEDYELAKMLTRKGSLFQND-
LAQRPKFTEEGYGSIKRVYVWTDEDKAIPPEFQLWQIENYKPDKVYRVQGGDHMLQLSKT-
NELAEILQEVADTYADLLAVAGGG

>HNL1_1st_8 contains 8 substitutions relative to HNL1
T11G,I12A,E79H,C81F,N104T,L152F,N156K,K236M
MATAHFVLIHGACHGAWIWYKLKPLLEAAGHKVTALDLAASGIDPRQIEQINTFDEYSEPLL-
TFMESLPQGEKVILVGHSFGGLNIALAADKYPEKISAAVFHTALMPDTEHS-
PSYVVDKFMEVFPDWKDTEFSTYTSNNETITGMKLGFKFMREKLYTNCPIEDYEL-
AKMLTRKGSFFQNDLAQRPKFTEEGYGSIKRVYVWTDEDKIFPPEFQLWQIENYKPDKVYR-
VQGGDHMLQLSKTNELAEILQEVADTYADLLAVAGGG

>HNL1_2nd_8 contains 8 substitutions relative to HNL1
I9V,H103L,K147L,E155N,T159Q,K175P,Q180I,L237P
MATAHFVLVHTICHGAWIWYKLKPLLEAAGHKVTALDLAASGIDPRQIEQINTFDEYSE-
PLLTFMESLPQGEKVILVGESCGGLNIALAADKYPEKISAAVFLNALMPDTEHSPSYVVD-
KFMEVFPDWKDTEFSTYTSNNETITGMLLGFKLMRNNLYQNCPIEDYELAKMLTRPGSFFIND-
LAQRPKFTEEGYGSIKRVYVWTDEDKIFPPEFQLWQIENYKPDKVYRVQGGDHKPQLSKT-
NELAEILQEVADTYADLLAVAGGG

>HNL1-16 contains 16 substitutions relative to HNL1
I9V,T11G,I12A,E79H,C81F,H103L,N104T,K147L,**L152F,E155N,N156K,T159Q,K175P,Q180I,K236M,L237P
MATAHFVLVHGACHGAWIWYKLKPLLEAAGHKVTALDLAASGIDPRQIEQINTFDEYSE-
PLLTFMESLPQGEKVILVGHSFGGLNIALAADKYPEKISAAVFLTALMPDTEHSPSYVVD-
KFMEVFPDWKDTEFSTYTSNNETITGMLLGFKFMRNKLYQNCPIEDYELAKMLTRPGSFFIND-

LAQRPKFTEEGYGSIKRVYVWTDEDKIFPPEFQLWQIENYKPDKVYRVQGGDHMPQLSKT-
NELAEILQEVADTYADLLAVAGGG

>HNL1_SigniSite_38;A4K,I9V,T11G,I12A,Q47K,L62M,T63E,F64L,G71N,E7
9H,C81F,A90M,D91E,H103L,N104T,T137G,I143L,K147L,L152F,E155N,N156
K,T159Q,Y166L,K175P,Q180I,R186A,I197V,D205N,Y222S,K223G,Y228M,V2
30I,G233A,K236M,L237P,T242P,Q250L,V252I
MATKHFVLVHGACHGAWIWYKLKPLLEAAGHKVTALDLAASGIDPRKIEQINTFDEYSE-
PLMELMESLPQNEKVILVGHSFGGLNIALAAEKYPEKISAAVFLTALMPDTEHSPSYVVD-
KFMEVFPDWKDTEFSTYGSNNETLTGMLLGFKFMRNKLYQNCPIEDLELAKMLTRPGSFFIND-
LAQAPKFTEEGYGSVKRVYVWTNEDKIFPPEFQLWQIENSGPDKVMRIQGADHMPQLSKP-
NELAEILLEIADTYADLLAVAGGG
